# CtBP1 coordinates synaptic, metabolic and contractile changes induced by denervation in skeletal muscle

**DOI:** 10.1101/2025.03.29.646097

**Authors:** Olivia Cattaneo, Gaetan Lopez, Jayasimman Rajendran, Florent Chabry, Alexandre Prola, Nicolas Liaudet, Sergei Startchik, Perrine Castets

**Author notes:** Contributed equally (Gaetan Lopez, Jayasimman Rajendran).

## Abstract

Nerve injury triggers dramatic atrophy of skeletal muscle, accompanied with synaptic and metabolic changes. Regulation of denervation-induced muscle fiber remodeling involves several factors governing genetic reprogramming and proteostasis changes. Here, we demonstrate that the transcriptional co-repressor CtBP1 coordinates synaptic and metabolic changes in muscle fibers upon denervation. CtBP1 was present both in sub- and non-synaptic myonuclei in innervated muscle. Although CtBP1 levels remained unchanged in denervated muscle, CtBP1 accumulated transiently in myonuclei after 2 days of denervation. *Ctbp1* knockdown perturbed the expression of a large set of activity-independent and -dependent genes in innervated and denervated skeletal muscles. CtBP1 loss had limited effect on the expression of most synaptic genes, but increased transcript levels of *Chrne,* encoding the adult ε sub-unit of acetylcholine receptors (AChR). However, it did not affect AChR turnover or maintenance of the post-synaptic compartment upon denervation. Importantly, we uncovered that *Ctbp1* knockdown promotes denervation-induced changes in metabolic gene expression, including most genes encoding proteins of the respiratory chain complexes. Consistently, it enhanced the switch towards slower, oxidative fibers in fast muscle after 2 weeks of denervation. Moreover, CtBP1 loss precipitated the profound ultrastructural remodeling of mitochondria network induced after denervation. Hence, our study unveils the role of CtBP1 in the integrated muscle response to denervation, with important implications for CtBP1-related muscle diseases.

**One-sentence summary:** Loss of CtBP1 perturbs synaptic, metabolic and contractile changes induced by denervation in skeletal muscle

## Introduction

Skeletal muscle contraction is controlled by motoneurons, which connect myofibers at the neuromuscular junction (NMJ). Efficient neurotransmission is ensured by the accumulation of synaptic proteins, such as acetylcholine receptors (AChRs) specifically in the highly specialized, sub-synaptic region of myofibers (*i.e.,* the endplate). Within each multinucleated myofiber, only the few myonuclei located at the endplate express synaptic genes, while synaptic gene expression is repressed in all other nuclei (*1, 2*). Nerve injury quickly triggers a switch back to a more embryonic genetic program in myonuclei, which includes the re-expression of the myogenic transcription factor myogenin, and in turn, of synaptic genes in non-synaptic regions of denervated myofibers (*3, 4*). Synaptic remodeling coordinates with long-term effects of denervation, which include severe muscle atrophy and a switch in the metabolic and contractile types of myofibers. The class II histone deacetylase (HDAC) 9 (*5*) and the transcriptional repressors Dach2 (*3*), Msy3 (*6*) and CtBP1 (*7*) have been involved in the repression of synaptic and myogenic gene expression in innervated muscle. In contrast, HDAC4 induction triggers denervation-induced release of their repression (*8–10*) and contributes to the up-regulation of atrophy-related genes and to the metabolic switch in denervated muscle (*10–12*). Whether other factors contribute to metabolic and synaptic remodeling upon denervation remains largely unknown.

C-terminal binding protein 1 (CtBP1) is a ubiquitous, evolutionary conserved transcriptional regulator forming complexes with DNA-binding proteins through a PXDLS motif (*13*). The *CTBP1* gene encodes a long isoform (CtBP1-L) and a short alternatively spliced isoform, called Brefeldin A-ADP ribosylated substrate (BARS or CtBP1-S). These two isoforms were first associated with transcriptional co-repressor activity and fission of Golgi tubular network, respectively (*14–16*). Whether the two isoforms exert both functions remains unclear. CtBP1 activity is regulated through post-translational modifications influencing its shuttling from cytoplasm to nuclei. In particular, p21-activated kinase 1 (Pak1), by phosphorylating CtBP1 on Ser158, sequesters it in the cytoplasm and inhibits its co-repressor activity, while favoring its fission activity (*17, 18*). Pak1-dependent CtBP1 nuclear export in skeletal muscle contributes to the release of myogenic and synaptic gene repression 3 days after denervation (*7*). Interestingly, mutations in *CTBP1* cause muscle dysfunction marked by dystrophic features and mitochondrial alterations as part of the *Hypotonia, Ataxia, Developmental Delay and Tooth-enamel* syndrome (HADDTS) (*19–21*). While the oncogenic role of CtBP1 has been widely reported (*22*), the pathomechanisms underlying HADDTS-associated muscle dysfunction are largely unknown.

Here we examined the physiological roles of CtBP1 in regulating activity-dependent processes in skeletal muscle. We show that nerve injury triggers a quick and transient accumulation of CtBP1 in myonuclei. Knocking down *Ctbp1* exacerbated denervation-induced changes in synaptic, contractile and metabolic gene expression, and accelerated the ultrastructural remodeling of mitochondria network. These findings indicate that CtBP1 coordinates the complex synaptic, contractile and metabolic changes occurring in skeletal muscle after denervation.

## Results

### CtBP1 expression is independent from neural activity

To get insights into the role of CtBP1 in muscle homeostasis, we first examined the expression pattern of CtBP1 isoforms in muscle cells. *Ctbp1-l* and *Ctbp1-s* transcripts, encoding CtBP1-L and CtBP1-S isoforms, arise from alternative transcription start sites (TSS – fig. S1A) and hence share all exons but exon 1 (Fig. 1A). RT-PCR confirmed the expression of both *Ctbp1-l* and *Ctbp1-s* transcripts in C2C12 mouse myoblasts and myotubes (Fig. 1B). Their transcript levels remained largely unchanged from myoblasts to 7-day-differentiated myotubes (Fig. 1C). RNAseq data available on *WashU Epigenome* Browser (15, 16) confirmed that levels of both transcripts are similar between undifferentiated and differentiated muscle cells in mice and humans (fig. S1A). Notably, *CTBP1-L* levels, relative to *CTBP1-S* levels, were much lower in humans than in mice (fig. S1A). At the protein level, CtBP1-S is 12 amino acids smaller than CtBP1-L, making the two isoforms undistinguishable by Western Blot. Total CtBP1 protein levels were unchanged between C2C12 myoblasts and myotubes, while Pak1 expression progressively decreased upon muscle cell differentiation (fig. S1B).

**Fig. 1.**
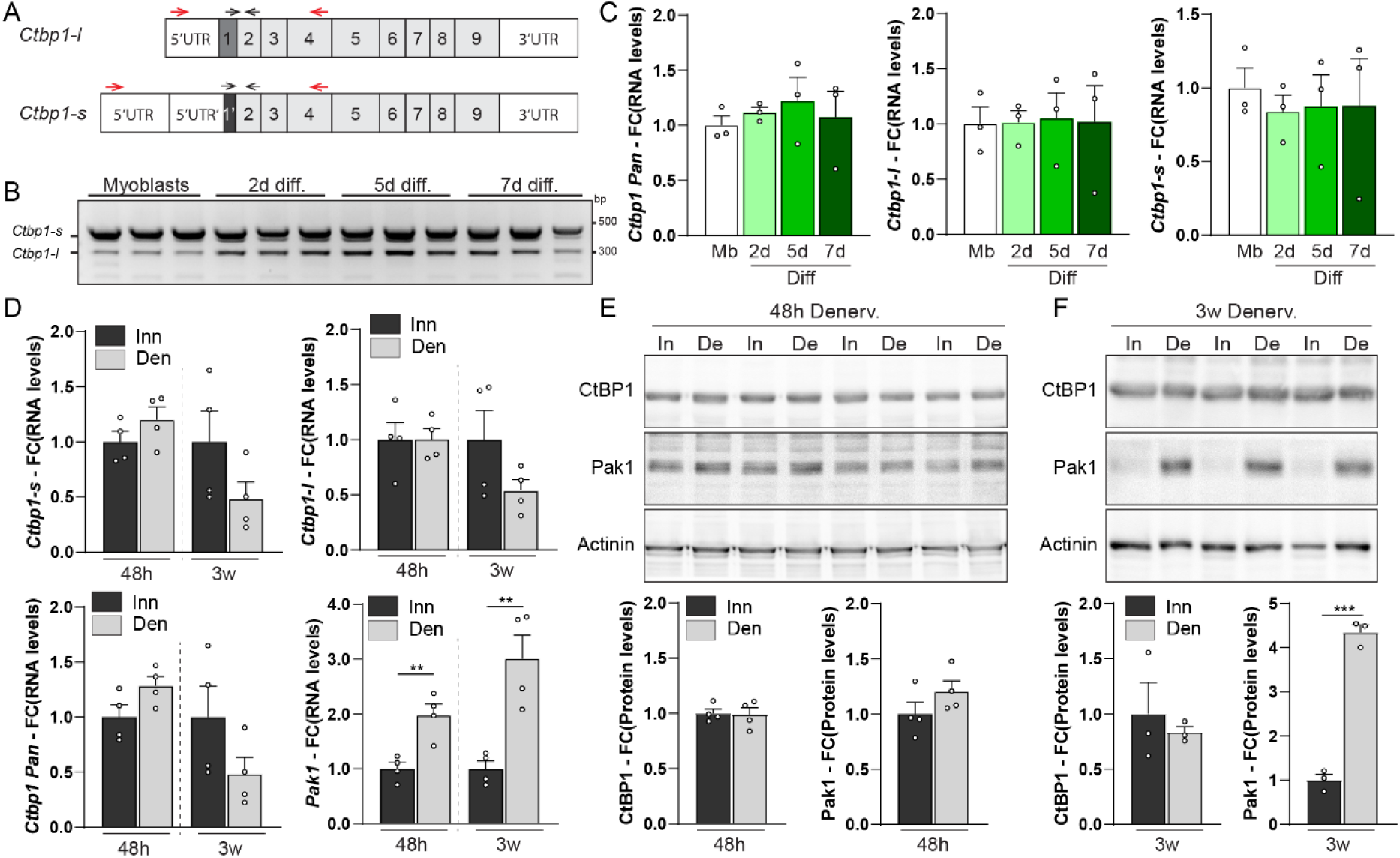
Expression of CtBP1-L and CtBP1-S remains unchanged upon nerve injury. **(A)** Organization of *Ctbp1-l* and *Ctbp1-s* mRNA in mouse. Exons and untranslated (UTR) sequences are represented by grey and white boxes, respectively. Primers used for PCR and qPCR are shown on top of mRNA with red and black arrows, respectively. (**B**) Expression of *Ctbp1-l* and *Ctbp1-s* is detected by PCR in C2C12 myoblasts and after 2 to 7 days (d) of differentiation (diff). n=3 independent samples. (**C**) Transcript levels of total *Ctbp1*, *Ctbp1-l* and *Ctbp1-s* in C2C12 myoblasts (Mb) and after 2 to 7 days (d) of differentiation (diff). Levels are relative to *Tbp* mRNA and to myoblast. (**D**) Transcript levels of total *Ctbp1*, *Ctbp1-l*, *Ctbp1-s* and *Pak1* in innervated (Inn) and denervated (Den – 48h and 3weeks) TA muscles. Levels are relative to *Tbp* mRNA and to innervated muscle. (**E** and **F**) Western Blot analysis of CtBP1 and Pak1 in innervated muscle (In) and 48h (E) or 3 weeks (F) after denervation (De). Protein levels are normalized to Actinin and to innervated muscle. All values are mean ± s.e.m.; n=3 (C) and 4 (D-F); ***p<0.001, Student’s t-test.

We next assessed the effect of neural activity on CtBP1 isoform expression pattern, by using sciatic nerve cut to trigger denervation of hind limb skeletal muscle. Total *Ctbp1* (*pan*), *Ctbp1-l* and *Ctbp1-s* transcript levels were unchanged in *tibialis anterior* (TA) 2 days and 3 weeks after inducing denervation, compared to innervated muscle (Fig. 1D). Total CtBP1 protein levels were also comparable in innervated and 16h- to 3-week-denervated TA muscles (Fig. 1, E and F, fig. S1C). In contrast, increase in *Pak1* transcript levels was detected in muscle 2 days after nerve injury (Fig. 1D) and the protein strongly accumulated after 7 days of denervation (Fig. 1, E and F, fig. S1C), as previously reported (*7*). Hence, although expression of the well-known CtBP1 regulator Pak1 depends on neural activity, expression of CtBP1 isoforms is unchanged upon nerve injury.

### CtBP1 accumulates in myonuclei shortly after denervation

A previous report links CtBP1 nuclear export with the release of synaptic gene repression in non-synaptic myonuclei 3 days after denervation (*7*). To get further insights into CtBP1 function in activity-dependent processes, we first examined its sub-cellular distribution in sub- and non-synaptic muscle regions. Staining of transversal muscle sections with CtBP1 antibody and α-bungarotoxin (Btx), which binds AChRs, showed a predominant accumulation of CtBP1 in sub- and non-synaptic nuclei, as well as at the NMJ (Fig. 2A). Immunostaining of isolated muscle fibers also gave a faint cytoplasmic dotty pattern of CtBP1, with a strong straining in myonuclei and at the NMJ (Fig. 2B). *Ctbp1* knockdown using adeno-associated virus isotype 9 (AAV) carrying small hairpin RNA (shRNA) injected in TA muscle strongly reduced the staining observed, confirming its specificity (fig. S2A). Using *stimulated emission depletion* (STED) microscopy, we showed that CtBP1 staining at the NMJ co-localizes predominantly with synaptophysin, a marker of the pre-synaptic compartment, with fainter dotty staining detected in the post-synaptic region (Fig. 2C). The pre-synaptic accumulation of CtBP1 is consistent with its known role in pre-synaptic active zones (*18, 23*). We next examined temporal changes in CtBP1 localization induced by denervation, by analyzing TA muscle 16h to 21 days after sciatic nerve cut. While CtBP1 distribution at 16h (fig. S2B) and 7 days (Fig. 2D) post-denervation was similar to innervated muscle, its nuclear accumulation increased 48h (Fig. 2D) and 21 days (fig. S2B) after nerve injury. These dynamic changes in CtBP1 localization were observed with two different antibodies (fig. S2C). To confirm these results, we stained single fibers isolated from denervated *extensor digitorum longus* (EDL) muscle with CtBP1 antibodies and Btx. Consistent with muscle section immunostaining, CtBP1 staining increased in non- and sub-synaptic myonuclei after 48h of denervation, compared to innervated fibers, with both antibodies (Fig. 2E and fig. S2D). After 7 days of denervation, CtBP1 nuclear staining normalized and appeared more diffuse around myonuclei (Fig. 2E and fig. S2E). Of note, CtBP1 staining at the NMJ was lost shortly after nerve injury (Fig. 2E), which may be caused by pre-synaptic vesicle exocytosis and the release of CtBP1 from active zones upon nerve cut. Together these results indicate that nerve injury triggers dynamic changes in CtBP1 sub-cellular localization in both sub- and non-synaptic regions of muscle fibers.

**Fig. 2.**
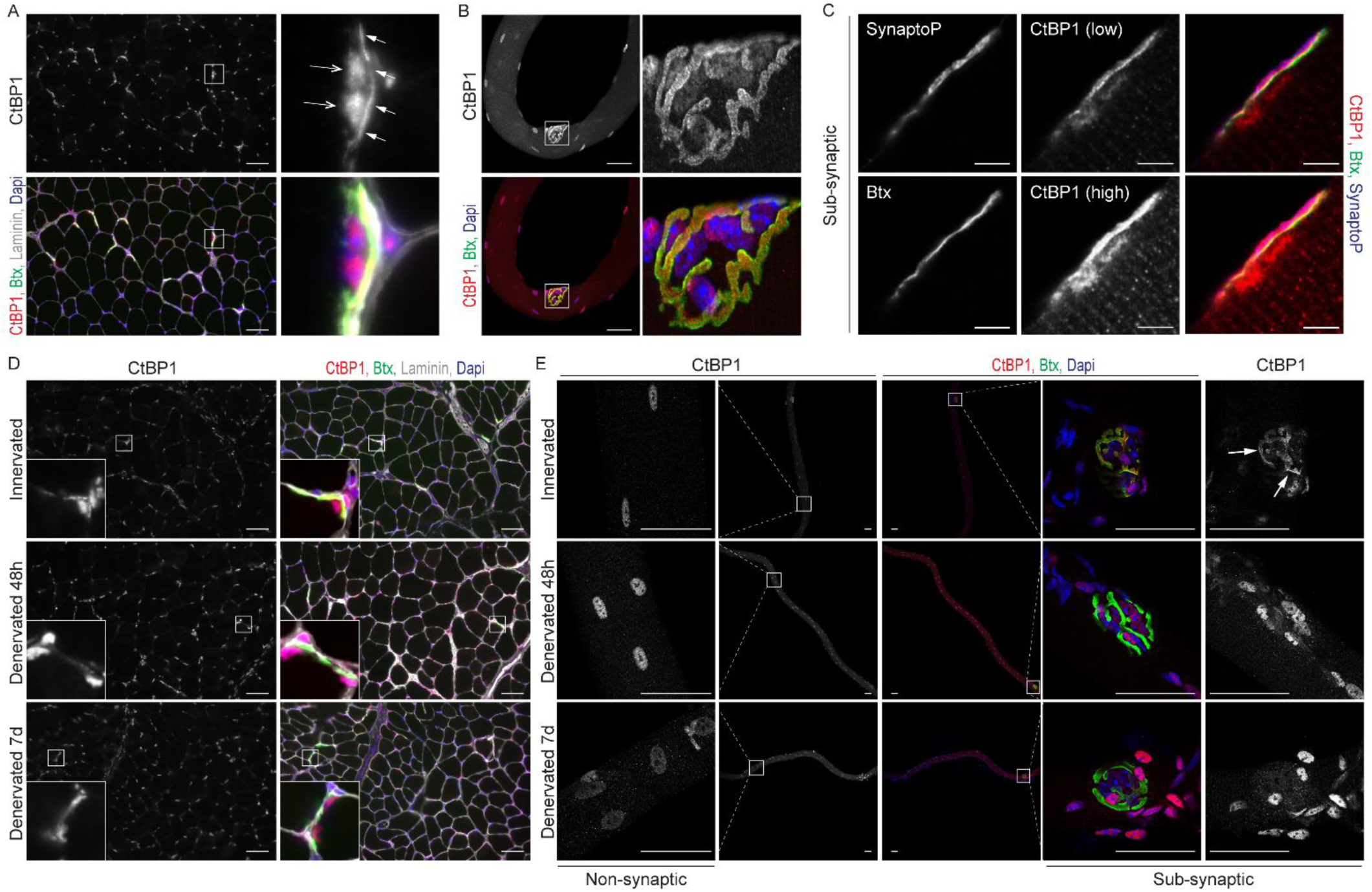
CtBP1 accumulates in myonuclei 48 h after denervation. (**A** and **B**) Staining of TA section (A) and EDL single fibers (B) with CtBP1 antibody and α-bungarotoxin (Btx) reveals accumulation of CtBP1 in non- and sub-(A, open arrows) myonuclei, as well as at the NMJ (A, arrows). Scale bar, 50µm. (**C**) STED microscopy of single fiber stained with CtBP1 and synaptophysin (SynaptoP) antibodies, and Btx, shows predominant CtBP1 accumulation in NMJ pre-synaptic compartment. Scale bar, 5μm. (**D** and **E**) Immunostaining of innervated and denervated (48h and 7 days) TA sections (D) and EDL single fibers (E) shows transient accumulation of CtBP1 in non- and sub-synaptic myonuclei, 48h after nerve injury. CtBP1 pre-synaptic staining (arrows, E) is lost in denervated muscle fibers. Scale bar, 50μm.

### *Ctbp1* knockdown has minor effect on histology of innervated and denervated muscles

To assess the functional consequences of CtBP1 nuclear-cytoplasmic trafficking in denervated muscle, we injected AAV9-shRNA directed against *Ctbp1* in both hind limb anterior compartments of 3-month-old mice to knockdown *Ctbp1* expression in TA and EDL muscles. Four weeks later, sciatic nerve was cut unilaterally to compare the effect of *Ctbp1* knockdown in innervated and denervated muscles (Fig. 3A). A single injection of AAV-sh*Ctbp1* was sufficient to abolish *Ctbp1-l* and *Ctbp1-s* transcript levels (Fig. 3B) and to strongly reduce total CtBP1 protein levels in TA muscle, compared to control muscle injected with scramble shRNA (Fig. 3, C and D). Of note, CtBP1 pre-synaptic accumulation was preserved in innervated muscle fibers treated with AAV-sh*Ctbp1*, which was consistent with a predominant effect of the AAV9-shRNA in muscle fibers (fig. S3A). As *Ctbp1* and *Ctbp2* transcripts share more than 70% sequence similarities, we evaluated *Ctbp2* expression by qPCR to assess the specificity of the AAV-shRNA and the lack of compensatory expression. Indeed, *Ctbp2* transcript levels were similar in TA muscles injected with AAV-sh*Ctbp1* or AAV-shScramble (Fig. 3E). Similarly, injection of AAV-shRNA directed against *Ctbp2* reduced specifically *Ctpb2* expression, while the co-injection of AAV-sh*Ctpb1* and AAV-sh*Ctpb2* decreased both *Ctbp1* and *Ctbp2* transcript levels (fig. S3, B and C).

**Fig. 3.**
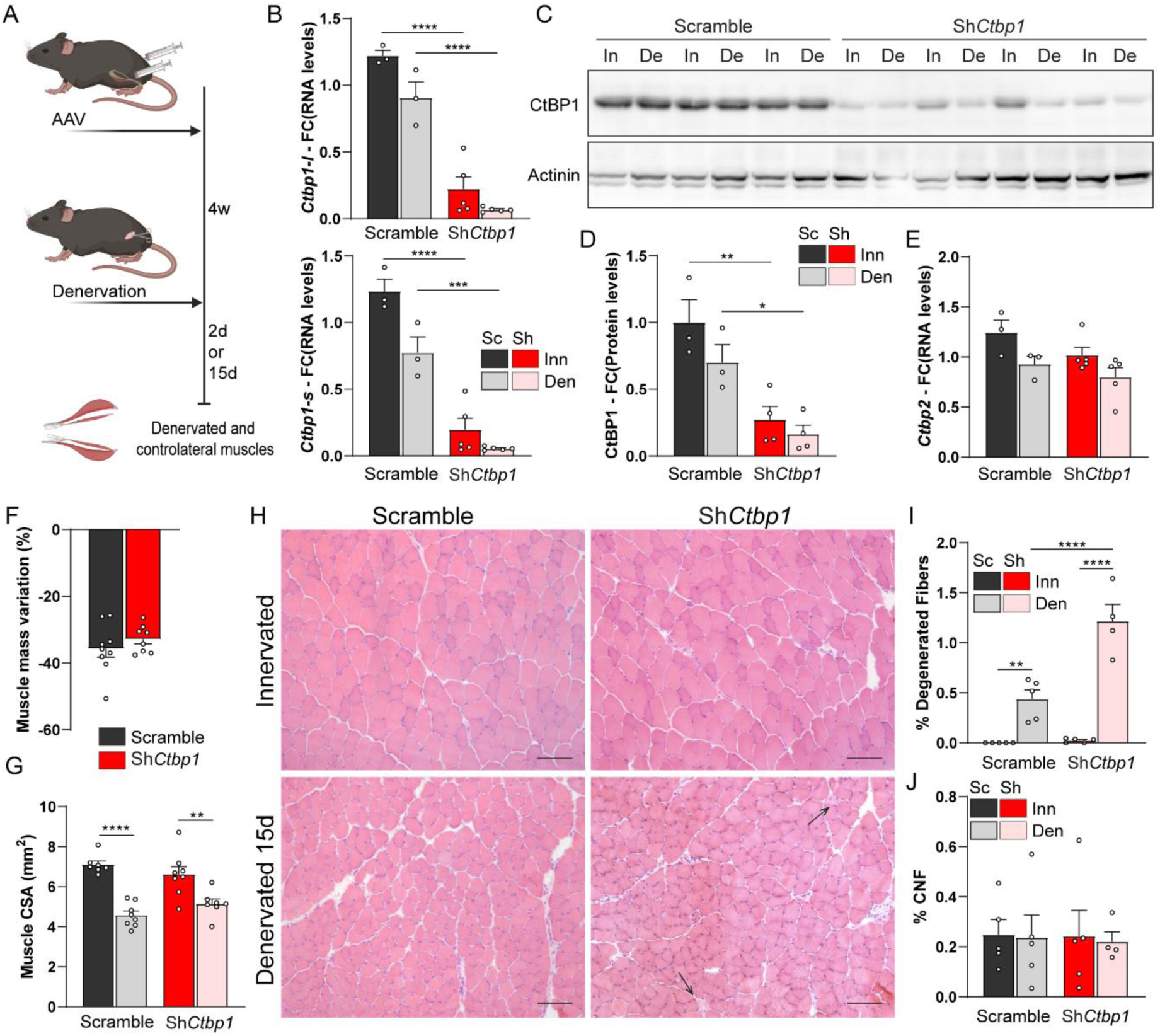
*Ctbp1* knockdown has minor effect on muscle histology. (**A**) Sciatic nerve was cut unilaterally 4 weeks after injecting AAV-sh*Ctbp1* or -shScramble, and TA/EDL muscles were analyzed 2 or 15 days later. (**B**) mRNA levels of *Ctbp1*-*l* and *Ctbp1-s* in innervated (Inn) and denervated (Den) TA muscles injected with AAV-sh*Ctbp1* (Sh) or -shScramble (Sc). Levels are relative to *Tbp* mRNA and to Scramble innervated. (**C** and **D**) Western Blot analysis of CtBP1 in innervated (In) and denervated (De) TA muscles injected with AAV-sh*Ctbp1* or -shScramble. Protein levels are normalized to actinin, and relative to Scramble innervated. Quantification of CtBP1 levels is given in (D). (**E**) mRNA levels of *Ctbp2* in innervated (Inn) and denervated (Den) TA muscles injected with AAV-sh*Ctbp1* (Sh) or - shScramble (Sc). Levels are relative to *Tbp* mRNA and to Scramble innervated. (**F**) Mass variation after 2 weeks of denervation for TA muscles injected with AAV-sh*Ctbp1* or -shScramble. (**G**) Cross sectional area (CSA) of innervated and denervated (2 weeks) TA muscles injected with AAV-sh*Ctbp1* or -shScramble. (**H**) H&E staining of innervated and 2-week-denervated TA muscles injected with AAV-sh*Ctbp1* or -shScramble. Scale bar, 100μm. (**I** and **J**) The proportion of degenerated fibers (I), but not of centronucleated fibers (CNF, J), is increased in denervated TA injected with AAV-sh*Ctbp1*, compared to Scramble. All values are mean ± s.e.m.; n=3Sc/5Sh (B, E), 3Sc/4Sh (C, D), 9Sc/8Sh (F), 7Sc/8Sh (G), 5 (I and J, except for Sh den, n=4); *p<0.05, **p<0.01, ***p<0.001, ****p<0.0001; two-way ANOVA with Tukey’s post-hoc.

Two weeks after denervation, the loss of muscle mass and reduction in muscle cross sectional area were similar between mice injected with AAV-shScramble or AAV-sh*Ctbp1* (Fig. 3, F and G). AAV-sh*Ctbp1* injection did not alter muscle histology in innervated conditions. It slightly increased the proportion of degenerating fibers after 2 weeks of denervation (Fig. 3, H and I), but the proportion of centronucleated fibers (*i.e.,* regenerating) remained unchanged compared to control TA (Fig. 3J). There was no major change in muscle mass loss or muscle histology after knocking down *Ctbp2* alone or both *Ctbp1* and *Ctbp2* (fig. S3, D-H). Hence, these results indicate that CtBP1 and/or CtBP2 loss has minor effect on muscle histology after denervation.

### *Ctbp1* knockdown does not perturb AChR dynamics and endplate maintenance

Previous reports establish that CtBP1 regulates synaptic vesicle exocytosis / recycling in pre-synaptic compartments (*18, 23*). As we detected a dotty staining for CtBP1 in the sub-synaptic region of muscle fibers, we examined whether it may correspond to vesicles enriched in newly formed or recycling AChRs. Consistent with increased AChR turnover upon denervation, Btx-positive dots in the endplate region, previously reported as endocytic vesicles containing AChRs (*24, 25*), accumulated further in single fibers isolated 7 days after denervation than in innervated fibers (Fig. 4A). Interestingly, some, but not all, Btx-positive vesicles co-localized with CtBP1 staining at the endplate (Fig. 4A). This suggests that CtBP1 may contribute to the regulation of post-synaptic vesicle trafficking. To assess the role of CtBP1 in AChR dynamics, we combined AAV-sh*Ctbp1* or AAV-shScramble injection with AChR turnover assay, using established procedures with sequential labelling of stable and newly formed AChRs with distinct fluorescent Btx (Fig. 4B) (*24, 26*). AAV-sh*Ctbp1*-infected fibers from innervated and 2-week-denervated TA muscles did not show major endplate alteration, as compared to AAV-shScramble muscle (Fig. 4C-E). In particular, CtBP1 deficiency altered neither endplate volume (Fig. 4C) nor endplate fragmentation (Fig. 4D) in TA innervated and denervated muscles, as compared to control muscle. Consistent with AChR destabilization and increased formation of new receptors in muscle after nerve injury, AChR turnover was strongly increased in control denervated muscles (Fig. 4, F and G). Similar increase in AChR turnover was observed after *Ctbp1* knockdown, suggesting that CtBP1 is dispensable for AChR dynamics (*e.g.,* internalization and recycling) in innervated and denervated conditions. These results suggest that denervation-induced changes in CtBP1 nuclear-cytoplasmic trafficking is not coupled with sub-synaptic regulation of AChR-containing vesicles.

**Fig. 4.**
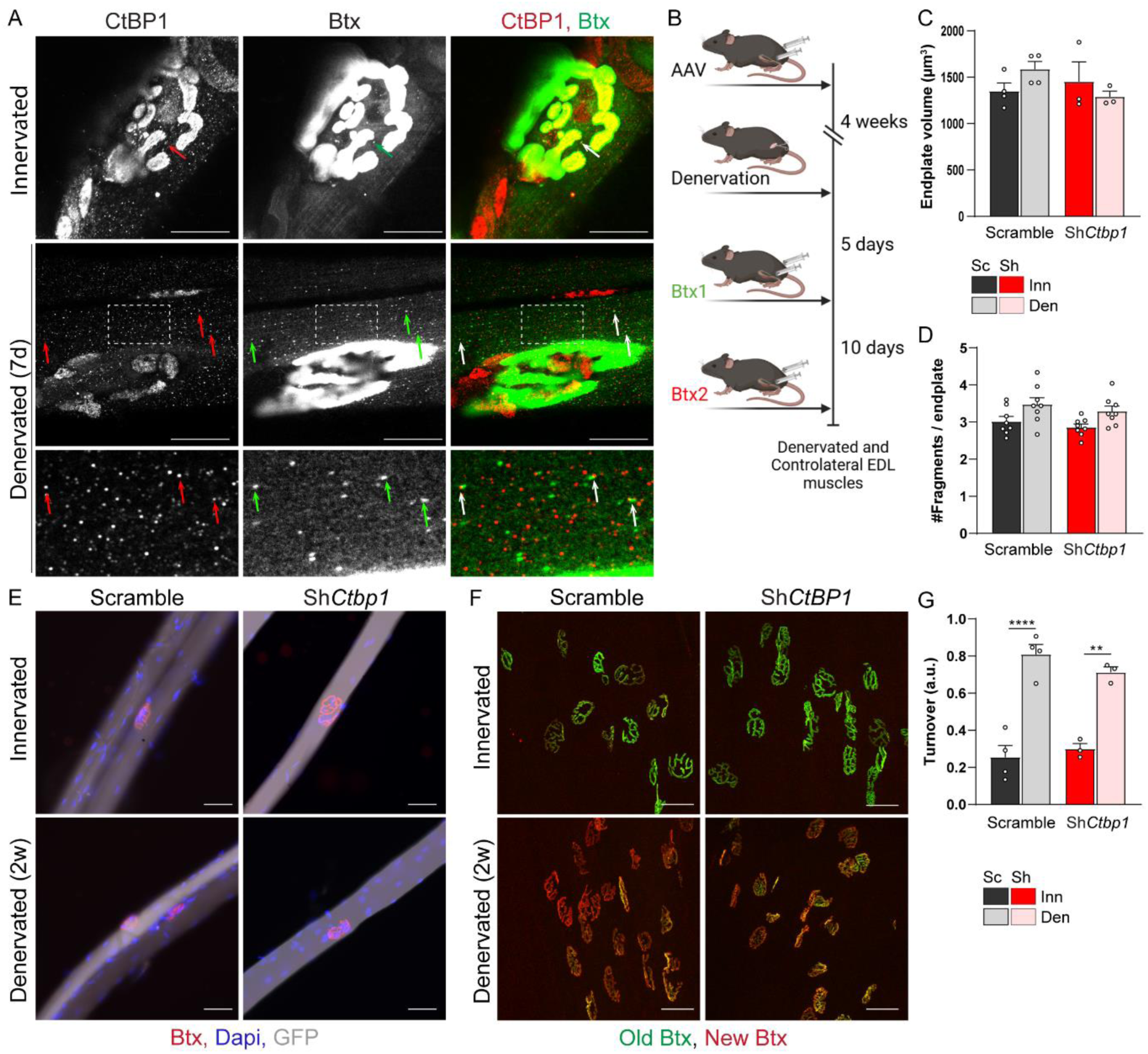
*Ctbp1* knockdown does not impair AChRs dynamics and endplate maintenance. (**A**) Immunostaining of permeabilized single EDL fibers with CtBP1 antibody and Btx shows that some AChR-containing vesicles stain positive for CtBP1 (arrows) in innervated and denervated muscle. Scale bar, 20 µm. (**B**) Injection of AAV-sh*Ctbp1* and -shScramble was combined with Btx injection to analyze AChR dynamics. (**C**-**E**) Btx staining of innervated (Inn) and denervated (Den) EDL fibers shows preserved endplate structure after AAV-sh*Ctbp1* (GFP-positive) infection (E). Quantification of endplate volume and fragmentation is given in C and D, respectively. Scale bar, 50 µm. (**F** and **G**) Whole-mount Btx staining of innervated and 2-week-denervated EDL injected with AAV-sh*Ctbp1* or - shScramble distinguishes “old” (green) and “new” (red) AChRs. Quantification of AChR turnover is given in G. Scale bar, 50 μm. All values are mean ± s.e.m.; n=8(D), 4Sc/3Sh (C, G); **p<0.01, ****p<0.0001; two-way ANOVA with Tukey’s post-hoc.

### CtBP1 regulates activity-dependent and -independent genes in skeletal muscle

CtBP1 transcriptional activity is associated with the repression of pro-apoptotic genes in cancer cells (*27, 28*), and of synaptic genes in innervated muscle (*7*). To test whether the transient accumulation of CtBP1 in myonuclei 2 days after denervation is associated with the regulation of specific activity-dependent genes, we performed RNAseq analysis of innervated and 2-day-denervated TA muscles injected with either AAV-shScramble or AAV-sh*Ctbp1*. The expression pattern of denervated and innervated muscles strongly differ, irrespective of CtBP1 modulation (fig. S4A). Comparison between innervated muscles injected with AAV-sh*Ctbp1* vs. -shScramble gave 566 differentially expressed (DE) genes, among which 53.0% were up-regulated upon *Ctbp1* knockdown (Fig. 5A). Similarly, 745 DE genes were identified in CtBP1-deficient denervated muscle compared to control denervated muscle, with 42.7% up-regulated genes. Interestingly, 48.7% of the genes deregulated with *Ctbp1* knockdown, were also DE between control innervated and denervated muscles (*i.e.,* activity-dependent) (Fig. 5A). Notably, 36.0 % correspond to known CtBP1 target genes (Fig. 5B), based on publicly available transcription factor databases (*29*). As recent evidence suggests that CtBP1 can also act as a transcriptional activator (*30, 31*), we considered genes up-regulated and down-regulated upon *Ctbp1* knockdown as direct or indirect targets repressed and induced by CtBP1, respectively. Activity-independent DE genes (hereafter referred to as X1 genes) included 53 genes deregulated in both innervated and denervated CtBP1-deficient muscles (Fig. 5, A and C). Other X1 genes were either deregulated in innervated or in denervated CtBP1-deficient muscle compared to control (fig. S4, B and C). Top X1 genes included *Ctbp1* as expected, as well as *Arrdc2*, *Sox7* and *Eppk1,* all three encoding factors involved in tumorigenesis and cell proliferation (Fig. 5C, fig. S4, B and C). Interestingly, among activity-dependent genes, 198 were only affected by *Ctbp1* knockdown in innervated muscle (hereafter referred to X2 genes, fig. S4D). For most of them, their deregulation mimicked the denervation effect (positive correlation with sh*Ctbp1*, X2^+^; Fig. 5D). Regulation of X2^+^ genes would hence fit with the model proposed for synaptic genes (*7*), in which denervation-induced changes in gene expression relies on the release of the repressive effect exerted by CtBP1 in innervated muscle (Fig. 5D). In contrast, X2^-^ genes (negative correlation with sh*Ctbp1*, fig. S4E), such as *Fgfbp1*, would be regulated similarly by CtBP1 in innervated muscle and by another independent factor after denervation (fig. S4E). In parallel, the deregulation of 330 activity-dependent genes only in denervated muscle was consistent with CtBP1 accumulation in myonuclei 2 days after denervation (fig. S4F). For most of these genes, the effect of denervation was dampened by CtBP1 activity (positive correlation with sh*Ctbp1*, X3^+^; Fig. 5E). Top X3^+^ genes included the metabolic genes *Gck*, *Ckmt2*, *Oxct1* and *Sdhb*, encoding glucokinase, mitochondrial creatine kinase, the ketolytic enzyme 3-oxoacid coA-transferase 1 and succinate deshydrogenase sub-unit B, respectively. These genes were down-regulated in denervated TA injected with AAV-shScramble, and even more with AAV-sh*Ctbp1*. Of note, *Tuft1*, which encodes a protein involved in oncogenesis and dental enamel mineralization (*32*) was the most up-regulated X3^+^ gene. Its deregulation downstream of CtBP1 may play a critical role in HADTTS. Inversely, for some X3 genes, the denervated effect may directly or indirectly rely on CtBP1 activity and its accumulation in nuclei (negative correlation with sh*Ctbp1*, X3^-^; Fig. 5F). X3^-^ genes included *Chrne* and *Tbx2*, encoding the adult ε sub-unit of AChR and TBX2, a known inhibitor of myogenesis (*33*). Finally, 53 activity-dependent genes were DE in both innervated and denervated CtBP1-deficient muscles compared to controls (referred to as X4 genes; Fig. 5G). Only *Chrna4, Scx, Myh6* and *Fhl2* showed an inverse regulation by sh*Ctbp1* in innervated and denervated muscles (Fig. 5G and fig. S4G). X4 genes included *Foxo3*, a known target repressed by CtBP1 (*34*), and *Chrna1*, encoding AChR sub-unit α, which were both up-regulated by denervation. To get further insight into the overall effect on *Ctbp1* knockdown on the muscle response to denervation, we used gene ontology (GO) enrichment analysis comparing AAV-sh*Ctbp1 vs.* shScramble and innervated *vs.* denervated muscles. As expected with the oncogenic role of CtBP1, pathways associated with cell division, cell movement and cytoskeletal were deregulated in CtBP1-deficient innervated muscle, compared to control muscle (Fig. 5H). Interestingly, *Ctbp1* knockdown led to a down-regulation of genes related to metabolic processes after nerve injury, as compared to control denervated muscle (Fig. 5H). In parallel, CtBP1 deficiency exacerbated the effect of denervation on the expression of genes related to cell cycle, nuclear regulation, RNA processing or (sub)cellular movement (Fig. 5H). Together, these results indicate that increased nuclear import of CtBP1 and its relocation on alternative target genes are required to release, limit or mediate the repression of specific synaptic and activity-dependent genes at early stages after denervation.

**Fig. 5.**
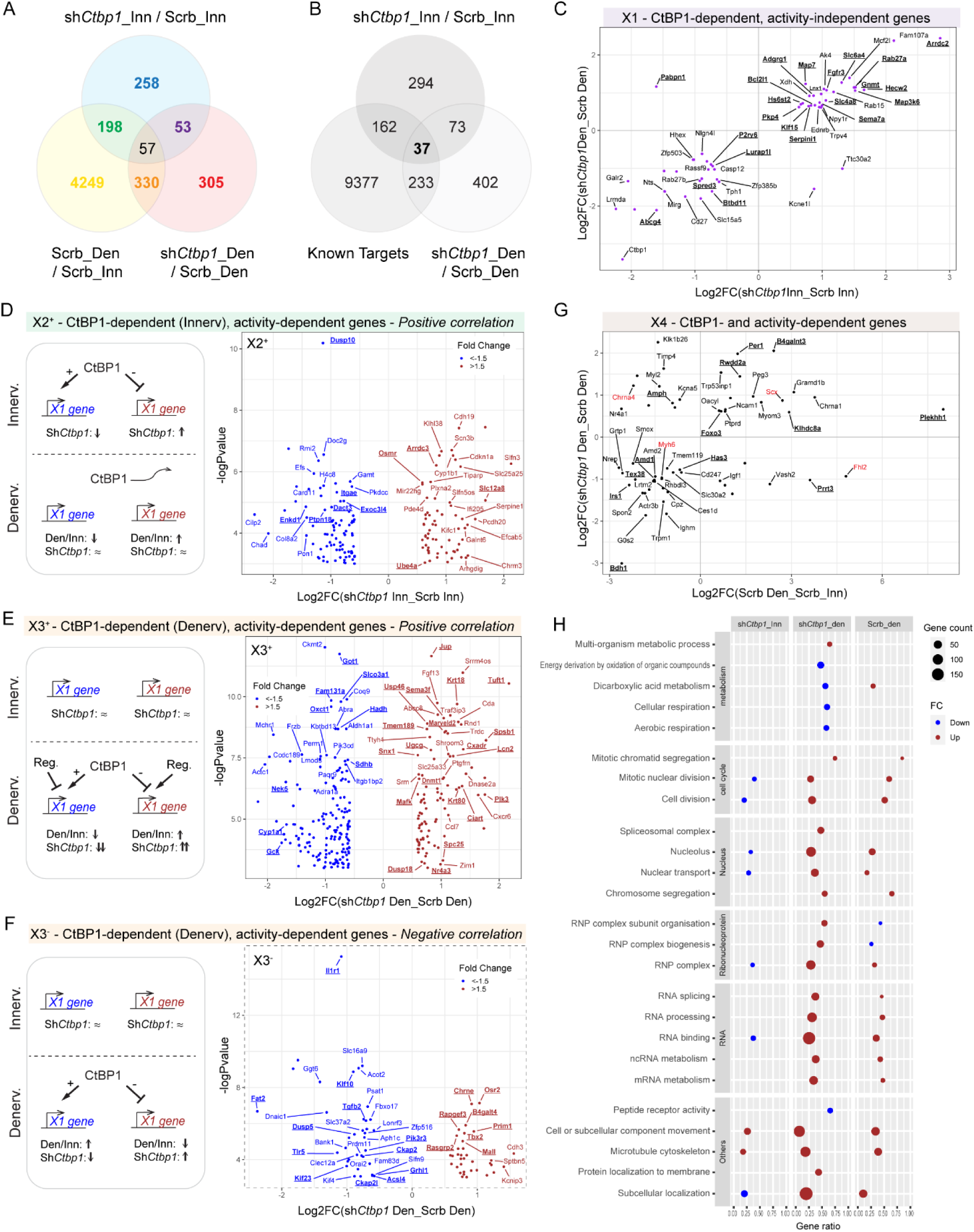
*Ctbp1* knockdown perturbs activity-dependent and -independent genes. (**A**) Venn diagram showing common DE genes between innervated (Inn) and denervated (Den) muscles infected with AAV-shScramble (Scrb; yellow), innervated muscles infected with AAV-sh*Ctbp1* or -Scrb (blue), and denervated muscles infected with AAV-sh*Ctbp1* or -Scrb (red). (**B**) Venn diagram showing common DE genes between CtBP1 known targets (extracted from *TFlink.net*), innervated muscles infected with AAV-sh*Ctbp1* or -Scrb (dark grey), and denervated muscles infected with AAV-sh*Ctbp1* or -Scrb (white). (**C**) Scatter plot showing activity-independent genes DE in AAV-sh*Ctbp1* innervated muscles (*vs*. AAV-Scrb innervated) and in AAV-sh*Ctbp1* denervated muscles (*vs*. AAV-Scrb denervated). Only some X1 genes show negative correlation between innervated and denervated changes. (**D**) *Ctbp1* knockdown has a similar effect on X2^+^ genes in innervated muscle as denervation. The volcano plot shows genes DE in AAV-sh*Ctbp1* innervated muscles (*vs*. AAV-Scrb innervated), with positive correlation with denervation-induced changes. (**E**) *Ctbp1* knockdown exacerbates the denervation effect on X3^+^ genes. The volcano plot shows genes DE in AAV-sh*Ctbp1* denervated muscles (*vs*. AAV-Scrb denervated), with positive correlation with denervation-induced changes. (**F**) *Ctbp1* knockdown limits the denervation effect on X3^-^ genes. The volcano plot shows genes DE in AAV-sh*Ctbp1* denervated muscles (*vs*. AAV-Scrb denervated), with negative correlation with denervation-induced changes. (**G**) Scatter plot showing activity-dependent genes DE in both AAV-sh*Ctbp1* innervated muscles (*vs*. AAV-Scrb innervated) and AAV-sh*Ctbp1* denervated muscles (*vs*. AAV-Scrb denervated). The plot shows the correlation between the effect of denervation and of AAV-sh*Ctbp1* in denervated muscle. Only some X4 genes (red) show an inverse correlation in innervated muscle (see fig. S5G). (**H**) GO enrichment analysis of genes DE after denervation and/or *Ctbp1* knockdown. Gene counts, up/down fold change, and gene ratio are indicated with the size, color and position of the bubble, respectively.

### CtBP1 regulates the expression of AChR α1 and ε sub-units

Up-regulation of synaptic genes after denervation relies on HDAC4 induction and the subsequent repression of *Dach2* and *Mitr/Hdac9*, which allows *Myog* expression (fig. S5A) (*3, 5, 8*). Release of CtBP1 repressive activity on *Myog* also contributes to *Myog* up-regulation upon denervation (*7*). As synaptic processes did not appear in GO analysis, we examined individual RNAseq data for genes encoding synaptic proteins or proteins involved in their regulation (Fig. 6A). As expected, expression of most synaptic (*e.g., Chrna1, Musk*) and myogenic (*e.g., Myog, Mef2d*) genes, as well as *Hdac4* and *Hdac6*, increased in denervated muscle injected with AAV-shScramble. Inversely, transcript levels of *Hdac9* and *Dach2*, as well as *Eno3* and *Pfkm*, encoding metabolic enzymes, decreased 2 days after denervation (Fig. 6A), as previously reported (*10*). Notably, transcript levels of *Chrne*, encoding the adult ε sub-unit of AChRs, were reduced in control muscle at 2 days post-denervation (Fig. 6A), while it was shown to be up-regulated at later stages (*2*). Using qPCR, we confirmed that *Ctbp1* knockdown increases *Chrna1* levels in innervated muscle, and further enhances its up-regulation in denervated muscle (Fig. 6B), as found with the RNAseq (X4 gene; Fig. 5G and Fig.6A). In contrast, expression of *Chrng, Myog* and *Hdac4* remained unchanged upon *Ctbp1* knockdown (Fig. 6, C-E). Knocking down *Ctbp2* or both *Ctbp1* and *Ctbp2* did not increase synaptic and myogenic gene expression in innervated muscle, neither (fig. S5B). As CtBP1 further accumulated in myonuclei 2 days after denervation, we examined genes known to be repressed upon denervation. Unexpectedly, *Dach2*, but not *Mitr*, was further repressed upon denervation when knocking down *Ctbp1* (Fig. 6, F and G). CtBP1 deficiency also further repressed the expression of *Eno3*, but not of *Pfkm*, two genes targeted by HDAC4 (*10*) (Fig. 6, H and I). More interestingly, we identified that *Ctbp1* knockdown increases *Chrne* transcript levels in innervated muscle and dampened its repression upon denervation (X3^-^ gene; Fig. 6J). This indicates that CtBP1 represses *Chrne* in innervated muscle, and contributes to its transient down-regulation 2 days after denervation, together with other repressors. These results indicate that temporal changes in CtBP1 localization contribute to the regulation of the expression of synaptic genes, including those encoding AChR sub-units α1 and ε in innervated muscle and after nerve injury.

**Fig. 6.**
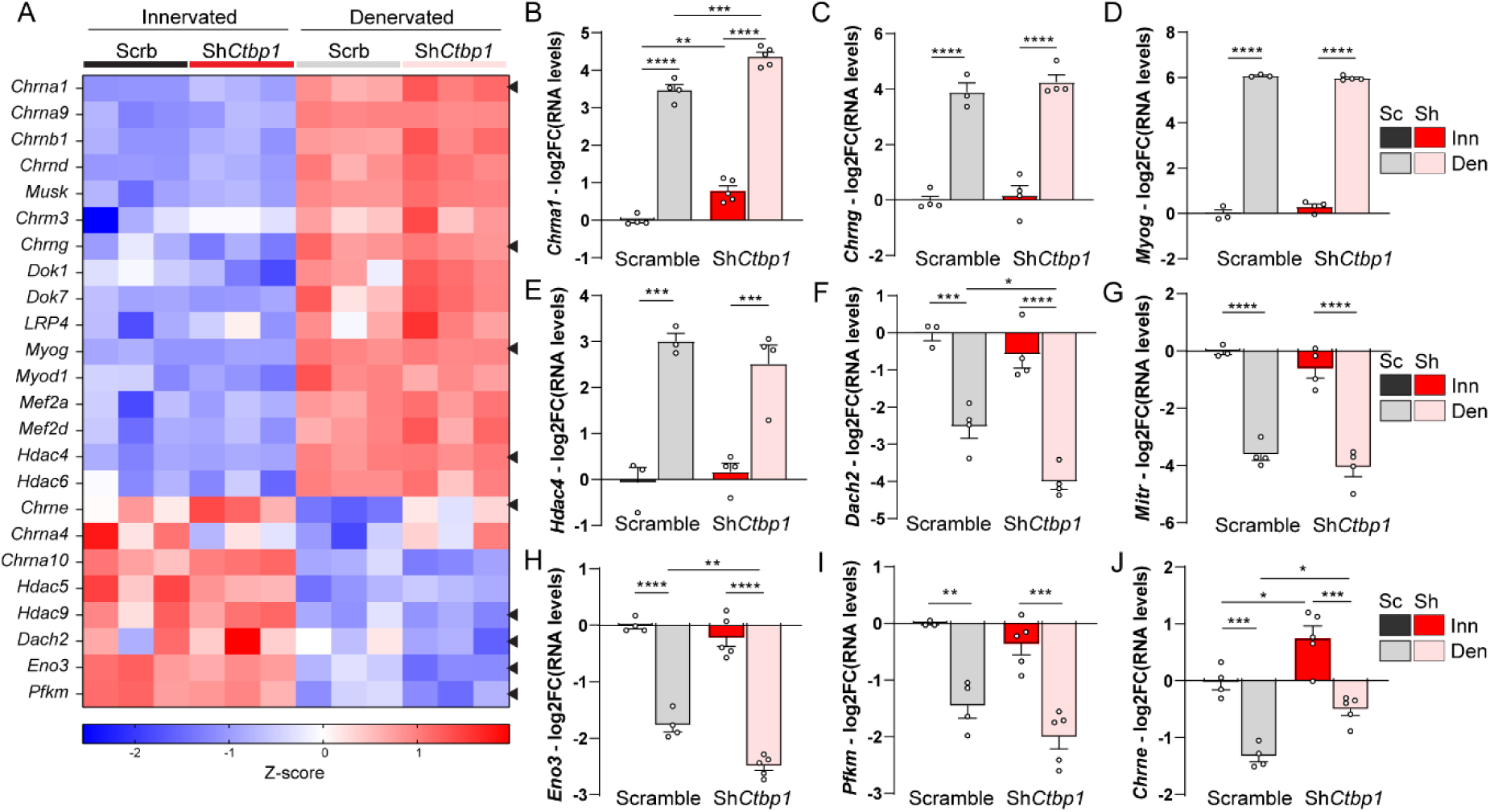
*Ctbp1* knockdown alters activity-dependent expression of *Chrna1* and *Chrne*. (**A**) Heatmap of z-scores computed based on log2FC of RNAseq counts for myogenic and synaptic genes in innervated and denervated, AAV-sh*Ctbp1* and -shScramble muscles. (**B-J**) mRNA levels of synaptic genes *Chrna1* (B), *Chrng* (C) and *Chrne* (J) and of the HDAC4 targets *Myog* (D), *Hdac4* (E), *Dach2* (F), *Mitr* (G), *Eno3* (H) and *Pfk* (I) after AAV-sh*CtBP1* or -shScramble injection in innervated or 48h-denervated TA muscles. Levels are relative to *Tbp* mRNA, normalized to AAV-shScramble innervated muscle, and analyzed as the log2 fold change (FC). Values are mean ± s.e.m.; n=4Sc/5Sh (B, H, J); 4ScInn/3ScDen/4Sh (C); 3Sc/4Sh (D, E); 3ScInn/4ScDen/4Sh (F, G); 3ScInn/4ScDen/5Sh (I); *p<0.05 **p<0.01 ***p<0.001 ****p<0.0001; two-way ANOVA with Tukey’s post-hoc.

### CtBP1 restrains fiber type switch but does not interfere with denervation-induced atrophy

Denervation triggers severe muscle atrophy, mainly driven by HDAC4 and FoxO pathways (*10, 35*). CtBPs were shown to repress FoxO3 activity and thereby to modulate the effect of the transcription factor ZEB1 on muscle atrophy (*36*). In TA muscle, denervation-induced muscle atrophy affects predominantly type IIB and IIX fibers, and is accompanied by a major shift towards type IIA oxidative fibers (*10, 24*). We first used RNAseq data to evaluate changes in the expression of genes encoding slow, fast or developmental isoforms of contractile proteins after knocking down *Ctbp1*. Denervation had overall a mild effect on contractile genes 2 days after nerve injury (Fig. 7A). Interestingly, *Ctbp1* knockdown repressed genes encoding fast isoforms of some contractile proteins (*e.g., Myh1* or *Myh4*) 2 days after denervation, and increased the expression of genes encoding slow isoforms of myosin light chain (*Myl*) or troponin (*Tpn*) in innervated muscle (Fig. 7A). We confirmed by qPCR that *Ctbp1* knockdown tends to repress *Myh4* (encoding myosin heavy chain (MHC) IIB) expression 2 days and 2 weeks after denervation (Fig. 7B). In parallel, *Myh2* (encoding MHCIIA) expression was strongly repressed 2 days after denervation, before being increased at 2 weeks (fig. S6A), as previously reported (*24*). However, *Ctbp1* knockdown did not alter *Myh2* transcript levels, compared to control muscle. In contrast, CtBP1 deficiency strongly increased the expression of *Myl2* (slow/cardiac isoform) in innervated muscle, and it limited its repression 2 days after denervation (Fig. 7C). As changes in contractile gene expression governs fiber type switch after nerve injury, we evaluated the proportion of type IIA/IIX/IIB fibers in innervated TA muscle and 2 weeks after denervation, based on immunostaining against MHCIIA/IIB (Fig. 7D). At 2 weeks, denervation increased the proportion of type IIA fibers in TA muscle, while the proportion of type IIX fibers was reduced (Fig. 7E). CtBP1-deficient innervated muscle showed an increased proportion of type IIA fibers (Fig. 7E), similar to that observed in control denervated muscle. Moreover, *Ctbp1* knockdown exacerbated the loss of type IIB fibers after denervation, as compared to control muscle (Fig. 7E). These results indicate that CtBP1 restrains the effect of denervation on fiber type shift, likely by regulating the transcriptional levels of contractile proteins.

**Fig. 7.**
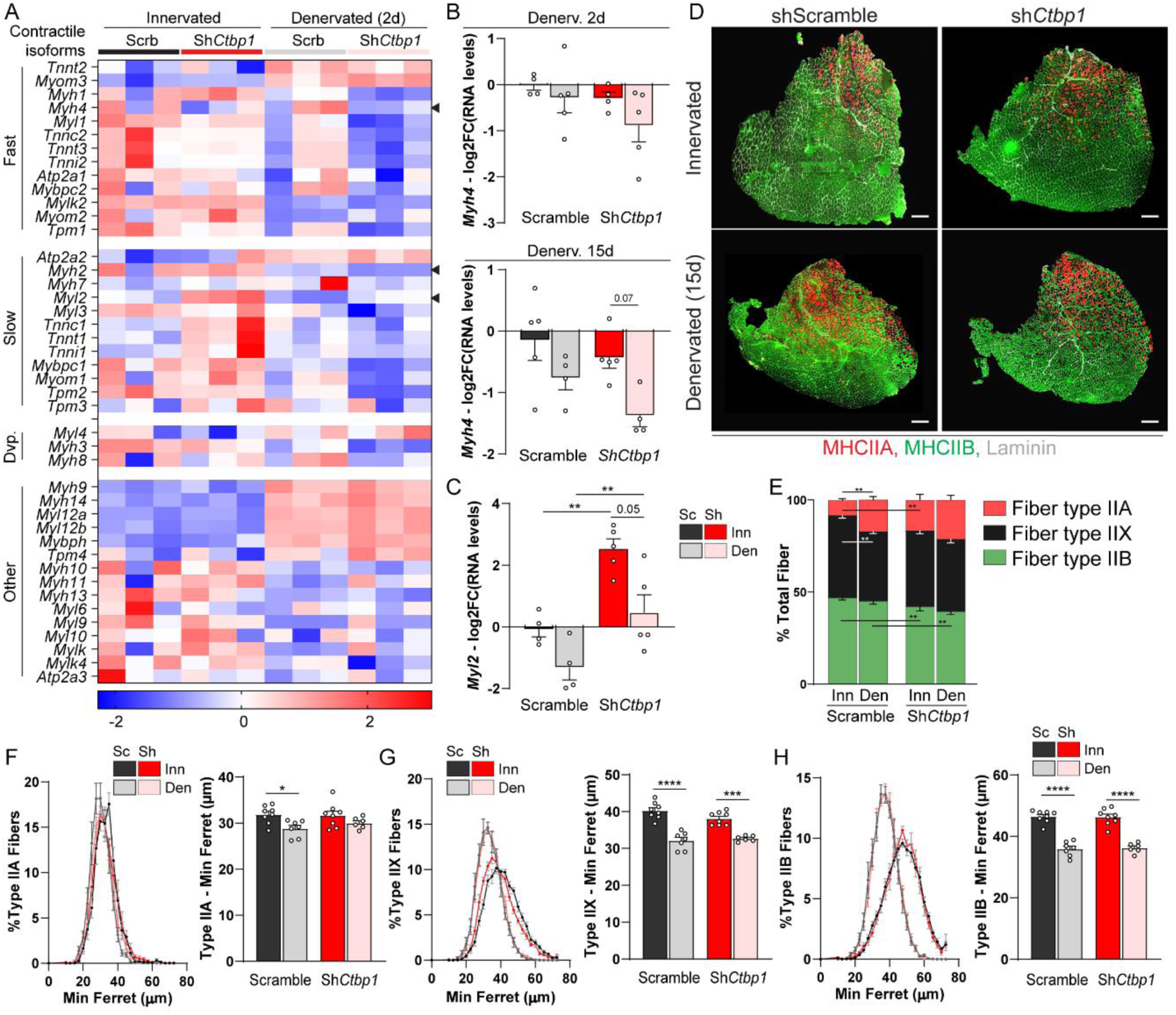
*Ctbp1* knockdown exacerbates the switch towards type IIA fibers. (**A**) Heatmap of z-scores computed based on log2FC of RNAseq counts for genes encoding fast, slow and developmental contractile protein isoforms in innervated and denervated, AAV-sh*Ctbp1* and -shScramble muscles. (**B** and **C**) mRNA levels of *Myh4* (B, 2 and 15 days) and *Myl2* (C, 2 days) after AAV-sh*CtBP1* (Sh) or - shScramble (Sc) injection in innervated (Inn) or denervated (Den) TA muscles. Levels are relative to *Tbp* mRNA, normalized to AAV-shScramble innervated muscle, and analyzed as the log2 fold change (FC). (**D** and **E**) Immunostaining for type IIA and IIB myosin heavy chains (MHC) and laminin, of innervated and 2-week-denervated TA muscles infected with AAV-sh*Ctbp1* or -shScramble. The proportion of types IIA, IIX and IIB fibers in TA muscle sections is given in E. Scale bar, 500µm. (**F**) Fiber size distribution and mean minimum ferret for types IIA, IIX and IIB fibers in innervated (Inn) and 2-week-denervated (Den) muscles infected with AAV-sh*Ctbp1* (Sh) or -shScramble (Sc). All values are mean ± s.e.m.; n=4Inn/5Den (B, 2d); 5Inn/4Den (B, 15d); 4Sc/5Sh (C); 8ScInn/7ScDen/8ShInn/6ShDen (E-H); *p<0.05, **p<0.01, ***p<0.001, ****p<0.0001; two-way ANOVA with Tukey’s post-hoc (B, E, F-G).

We next evaluated the effect of knocking down *Ctbp1* on muscle atrophy. RNASeq data confirmed that genes encoding proteins involved in catabolic processes, *i.e.,* proteasome machinery, calpains, autophagic components, were strongly up-regulated 2 days after denervation (fig. S6B). However, their expression was largely unchanged between control and CtBP1-deficient muscles, including for the atrogenes *Fbxo32* and *Trim63*, which are directly regulated by FoxO3. Only *Fbxo40* and *Nedd4,* which encode two E3 ubiquitin ligases associated with denervation-induced muscle atrophy (*37, 38*), showed reduced levels with *Ctbp1* knockdown in denervated muscle (fig. S6B). By qPCR, we confirmed that *Nedd4* expression is repressed in innervated muscle injected with AAV-sh*Ctbp1*, with a limited up-regulation after 2 weeks of denervation, as compared to control muscle (fig. S6C). In contrast, there was no major change in *Fbxo40* levels between mutant and control muscle 2 weeks after denervation (fig. S6C). These results indicate that *Ctbp1* knockdown has minor effect on atrophy-related gene expression in innervated muscle and after denervation. Consistently, *Ctbp1* knockdown did not alter fiber size, irrespective of the fiber type, in innervated muscle (Fig. 7F-H). Furthermore, it did not prevent or exacerbate the reduction in fiber size induced by denervation (Fig. 7F-H). These results demonstrate that CtBP1 restrains contractile fiber type switch induced by denervation, without interfering with atrophying processes.

### CtBP1 limits the oxidative shift and mitochondrial remodeling after denervation

GO analysis suggests a major role of CtBP1 in regulating metabolic paths, including cell respiration, in denervated muscle (Fig. 5H). Although denervation induces a switch from glycolytic to oxidative fibers in TA muscle, its effect on the different metabolic processes remains unclear. We showed that denervation perturbs the expression of genes associated with glycolysis (fig. S7A) and tricarboxylic acid cycle (TCA; fig. S7B), although with a mixed effect, with genes up- and down-regulated in each path in 2-day-denervated muscle. In contrast, nuclear genes encoding sub-units of the respiratory chain (complex I to V) were overall repressed 2 days after denervation (Fig.8A and fig. S7C). Importantly, repression of these metabolic genes was enhanced by *Ctbp1* knockdown, indicating that CtBP1 antagonizes the metabolic effect of denervation (X3^+^ genes; Fig. 8A and fig. S7A-C). Two weeks after denervation, expression of *Ndufs1, Ndufs2, Ndufv1* and *Sdhb* (encoding proteins of the respiratory chain) normalized in control muscle compared to innervated muscle (Fig. 8B and fig. S8A). Only *Cox7b* levels remained lower in 2-week-denervated muscle (Fig. 8B). At this stage, the repressive effect of AAV-sh*Ctbp1* on these metabolic genes was still detected, as compared to control muscle (Fig. 8B and fig. S8A). However, despite these transcriptional changes, protein levels of complex I to V of the respiratory chain remained similar after denervation or *Ctbp1* knockdown, 2 days and 2 weeks after denervation (fig. S8, B-E). This discrepancy may involve low turnover rate for complexed proteins of the respiratory chain. To test whether CtBP1 deficiency has an impact on respiratory chain function, we assessed complex I activity by performing NADH staining on TA muscle sections (Fig. 8C). As expected with the oxidative shift of TA muscle after nerve injury, the proportion of fibers with low NADH staining decreased 2 weeks after denervation in control muscle (Fig. 8D). Interestingly, *Ctbp1* knockdown exacerbated this switch, with a reduced proportion of fibers with low NADH staining in innervated muscle, and an increased proportion of fibers with high NADH staining in both innervated and denervated muscles, compared to controls (Fig. 8D). This indicates that CtBP1 deficiency increases complex I activity and precipitates the oxidative switch occurring after denervation. To go further, we examined mitochondria network in innervated and denervated muscles using electronic microscopy. Using longitudinal sections, we identified that mitochondria size increases in control muscle 2 weeks after nerve injury, leading to elongated structures in between sarcomeres (fig. S9A). Consistently, mitochondria area increased, while their mean circularity decreased after denervation (fig. S9, B and C). Interestingly, *Ctbp1* knockdown dampened denervation-induced changes in mitochondria size, leading to reduced mitochondria area in denervated muscle, as compared to control denervated muscle (fig. S9, A-C). To further characterize mitochondria remodeling upon denervation, we immunostained isolated fibers for TOMM20, a marker of the outer mitochondrial membrane (Fig. 8E). Using 3D analysis, we uncovered that the mitochondria network shifts from a transversal orientation compared to fibers, to a longitudinal orientation after 2 and even more so 3 weeks of denervation (Fig. 8E and fig. S9D). Using MATLAB software, we showed that the proportion of mitochondria forming an angle of 60-90° with the fiber decreased from 88.5% to 18.7% 3 weeks after denervation (Fig. 8F). This shift was also observed in muscle injected with AAV-sh*Ctpb1*, indicating that the reduced mitochondria size observed on longitudinal sections does not correspond to a defective re-orientation of the network. We thus evaluated changes in mitochondria volume using Imaris 3D volumetric reconstitution (fig. S9D). In both control and CtBP1-deficient muscles, the median mitochondria volume decreased after 2 and 3 weeks of denervation, compared to innervated muscle (fig. S9E). Consistently, the frequency distribution of mitochondria volumes was shifted to smaller mitochondria in control muscle after 2 and 3 weeks of denervation (Fig. 8G). Interestingly, CtBP1 deficiency already shifted the frequency distribution to smaller mitochondria in innervated muscle, and the distribution at 2 weeks of denervation was similar to the distribution observed at 3 weeks of denervation in control muscle (Fig. 9G). Hence, CtBP1 deficiency accelerates mitochondria size reduction triggered by denervation and occurring with the re-orientation of the network. Of note, this remodeling occurred without any changes in the transcript levels of genes encoding mitochondria fusion/fission machinery (*e.g., Mfn2* or *Opa1*) 2 days or 2 weeks after nerve injury, or after *Ctbp1* knockdown (fig. S9, F and G). Together, these data demonstrate that CtBP1 regulates metabolic changes in muscle after nerve injury, by limiting metabolic gene repression and mitochondria remodeling.

**Fig. 8.**
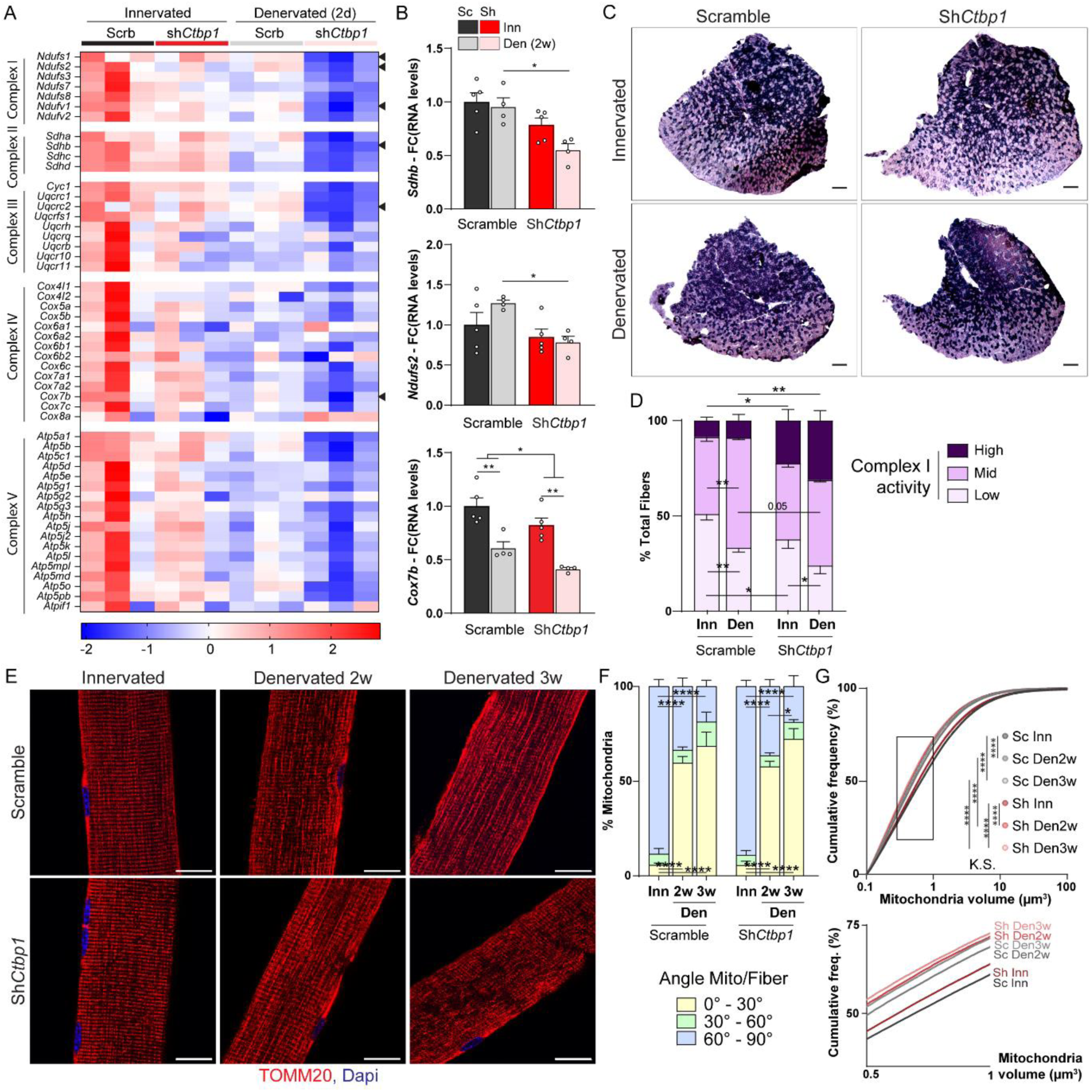
*Ctbp1* knockdown alters mitochondria activity and remodeling. (**A**) Heatmap of z-scores computed based on log2FC of RNAseq counts for nuclear genes encoding proteins of respiratory chain complexes I to V, in innervated and 2-day-denervated, AAV-sh*Ctbp1* and -shScramble muscles. (**B**) mRNA levels of genes encoding components of the respiratory complex I (*Ndufs2*), II (*Sdhb*) and IV (*Cox7b*) after AAV-sh*Ctbp1* (Sh) or -shScramble (Sc) injection in innervated (Inn) and 2-week-denervated (Den) TA muscles. Levels are relative to *Tbp* mRNA and normalized to AAV-shScramble innervated muscle. (**C** and **D**) NADH staining of innervated and 2-week-denervated TA muscles injected with AAV-sh*Ctbp1* or -shScramble. The proportion of fibers with high-, mid- and low-intensity staining is given in D. Scale bar, 500μm. (**E-G**) TOMM20 immunostaining of innervated and 2-/3-week-denervated TA muscles injected with AAV-sh*Ctbp1* or -shScramble shows mitochondria reorientation in denervated muscles. The proportion of mitochondria showing an angle of 0-30°, 30-60° or 60-90° as compared to the fiber is given in F. The cumulative frequency of mitochondria according to their volume is given in G. The 3D reconstitution of the pictures is given in fig. S9D. Scale bar, 20 µm. All values are mean ± s.e.m.; n=5Inn/4Den (B-D), 5 (E-G); *p<0.05, **p<0.01, ****p<0.0001; two-way ANOVA with Tukey’s post-hoc (B, D, F), Kolmogorov-Smirnov’s test (G).

## Discussion

Nerve injury triggers major synaptic, metabolic and contractile changes in muscle fibers, which are tightly regulated with time. Although different factors have been shown to contribute to this complex remodeling of denervated fibers, the integrated mechanisms underlying the muscle response to nerve injury is not fully understood. Here, we establish that dynamic activity-dependent nuclear-cytoplasmic trafficking of CtBP1 is essential for the regulation of synaptic and metabolic gene expression and that CtBP1 prevents excessive metabolic and contractile shift after denervation.

The vast majority of CtBP1 studies concerns its oncogenic role via transcriptional regulation of apoptosis, inflammation and cell cycle (*22*). In the neuromuscular system, CtBP1 is linked to the repression of myogenic and synaptic genes in mature muscle fibers (*7*), as well as with pre-synaptic vesicle recycling (*18, 23*). We here showed that CtBP1 is present in both sub- and non-synaptic myonuclei. Moreover, its nuclear accumulation transiently increased 48h after denervation in both types of myonuclei, independently from changes in its protein levels. This temporal regulation suggests that CtBP1 exerts activity-dependent functions in myonuclei and/or cytoplasm. Protein levels of Pak1, which drives CtBP1 nuclear export 3 days after denervation (*7*), were not yet changed 48h after nerve injury. Different post-translational modifications regulate CtBP1 nuclear-cytoplasmic trafficking in non-muscle cells. Especially, small ubiquitin-like modifier (SUMO) conjugation (sumoylation) of CtBP1, involving PIAS, RanBP2 or Cbx4 E3 ubiquitin ligases, increases its nuclear import (*39–41*). HDAC4 nuclear import, which also increases shortly after denervation, is promoted as well by RanBP2-dependent sumoylation (*42*). Inversely, activity of the myogenic transcription factors MEF2, which is tightly regulated after nerve injury, is inhibited by sumoylation (*43, 44*). Whether sumoylation changes contribute to CtPB1 nuclear-cytoplasmic trafficking in skeletal muscle requires further investigation.

The nuclear accumulation of CtBP1 2 days after nerve injury raised the question on the genes targeted at this stage. Knockdown assay confirmed that CtBP1 contributes to the repression of *Chrna1* in innervated muscle, while minor effect was found for *Myog* and *Chrng*. The use of AAV prevented muscle necrosis, which could account for the observed discrepancy in *Myog* expression changes compared to the results reported by Thomas et al., who employed muscle electroporation (*7*). *Chrna1* up-regulation was also enhanced after denervation in CtBP1-deficient muscle, while *Myog* expression was similarly increased. By forming repressive complex with MITR, HDAC7 and/or ZEB1, CtBP1 may directly inhibit MEF2 factors at E-boxes of some synaptic gene promoters and thereby repress their expression in non-synaptic muscle regions, independently from myogenin regulation (*45–47*). We further establish that CtBP1 plays a critical role in the regulation of *Chrne* expression. The selective expression of *Chrne* in sub-synaptic myonuclei, even after denervation, is linked to permissive effectors, such as p300 and ETV5/GABP, acting at the NMJ (*48, 49*). We identified that *Chrne* expression decreases quickly after denervation, before increasing as described before (*2*). *Ctbp1* knockdown increased *Chrne* expression in innervated muscle and limited its repression after denervation. CtBP1 may hence directly inhibit *Chrne* expression by forming repressive complex with HDAC1 and/or inhibiting p300 activity at ETS-binding sites of *Chrne* promoter in non- and sub-synaptic regions, respectively (*48, 50, 51*). Especially, *Chrne* repression may rely on CtBP1 transient import in sub-synaptic myonuclei shortly after nerve injury. Other repressors may contribute to *Chrne* repression at this stage, as *Ctbp1* knockdown did not abolish the repressive effect of denervation on the gene.

In neuron, CtBP1 traffics between nuclei and pre-synaptic compartments, where it regulates genes involved in synaptogenesis and synaptic vesicle recycling, respectively (*18, 23*). Its localization is tightly regulated by neural activity and relies on CtBP1interaction with Bassoon/Piccolo in active zones.

At the NMJ, we identified that pre-synaptic CtBP1 is lost quickly after axonal injury. Nerve injury may provoke massive release of pre-synaptic vesicles and/or local CtBP1 degradation during Wallerian degeneration. By analogy with neurons, we hypothesized that CtBP1 nuclear export detected 7 days after denervation may associate with synaptic functions. CtBP1 was present in some internalized AChR-containing vesicles at the endplate. However, *Ctbp1* knockdown did not affect endplate maintenance or AChR turnover in innervated and denervated muscles. This suggests that CtBP1 is dispensable for the regulation of post-synaptic vesicle recycling and trafficking.

In parallel with synaptic remodeling, we uncover that CtBP1 plays a key role in the regulation of metabolic and contractile properties of skeletal muscle after denervation. *Ctbp1* knockdown exacerbated the repression of genes related to glycolysis and mitochondrial respiration at 2 days of denervation. This result indicates that CtBP1 nuclear import is required to restrain metabolic gene repression upon nerve injury. CtBP1 may act as a transcriptional activator on these genes, as described in some inflammatory responses when complexed with p300 (*30, 31*), or it may indirectly promote their expression by repressing the expression or inhibiting the activity of specific transcriptional repressors. In several cell types, CtBP1 acts as a metabolic sensor by binding NADH, and to a lower extent NAD^+^ (*52–54*). NADH binding regulates the nuclear import, the oligomerization and the transcriptional activity of CtBP1 (*55–59*). Interactions of CtBP1 with transcriptional co-regulators are either increased or released upon NADH binding, leading to distinct transcriptional consequences upon changes in NADH/NAD^+^ levels (*59–61*). Interestingly, denervation confers transient insulin resistance to skeletal muscle 1-3 days after nerve injury (*62, 63*). This complex effect may involve changes in Akt activity, as well as opposite changes in *Glut1/Glut4* expression (*64–66*). By perturbing glycolytic flux and thereby free NADH levels, insulin resistance may trigger changes in CtBP1 activity shortly after denervation. Especially, this may release CtBP1 activity on some genes 2 days after denervation (*e.g., Chrna1*), while reinforcing its repressing or permissive activity on others (*e.g.,* metabolic genes). Despite its repressive effect on metabolic genes at early denervation stage, *Ctbp1* knockdown increased activity of respiratory chain complex I after prolonged denervation and thereby exacerbated the effect of nerve injury. The switch towards slower oxidative fibers in fast TA muscle after denervation involves the combined induction of HDAC4 (*10*), PGC1α (*67*) and myogenin (*68*). In this context, changes in mitochondria network have been debated (*69–76*). We here report that prolonged denervation triggers profound mitochondria remodeling with a re-orientation of the network and a reduction in mitochondria volume. These changes likely involve finely-tuned regulation of mitochondria fusion/fission, and may reflect the switch to oxidative fibers in TA muscle and/or the reinduction of the embryonic program after denervation (*77*). *Ctbp1* knockdown exacerbated the oxidative switch of TA muscle and the reduction in mitochondria volume induced by denervation. In contrast with its role in Golgi and vesicle membrane fission, CtBP1, and more specifically CtBP1-S, may favor mitochondria fusion, by regulating lipid organization and/or fusion/fission machinery components. Importantly, two reports suggest that mutations in *CTBP1* gene cause mitochondria dysfunction with reduced complex I and IV activity in muscle from HADDTS patients (*20, 21, 27*). The two corresponding mutations (*i.e.,* p.Arg331Trp and p.Gln439ValfsTer84) are located in the PXDLS domain of CtBP1 and may alter its interactions with DNA-binding proteins. Abnormal repression of metabolic genes and mitochondria fission/fragmentation, as observed with *Ctbp1* knockdown, may contribute to mitochondria dysfunction in patients. Muscle biopsies from HADDTS patients also display predominance of slow fiber types, which was consistent with the exacerbated switch towards type IIA fibers in CtBP1-deficient TA muscle. Accordingly, we showed that CtBP1 regulates genes encoding fast and slow contractile protein isoforms. Finally, despite the reported role of CtBP1 in repressing FoxO3 activity (*34*), CtBP1 deficiency did not perturb denervation-induced muscle atrophy and had no major effect on atrophy-related gene expression.

In conclusion, our work identifies an important role of CtBP1 in muscle homeostasis, by coordinating the regulation of contractile, metabolic and synaptic remodeling after denervation. CtBP1 deregulation may hence contribute to the complex pathogenic effects leading to muscle dysfunction in neuromuscular diseases and sarcopenia.

## Materials and Methods

### Animals

C57BL6 control mice were kept in a conventional animal facility with a fixed 12h dark/light cycle at 23°C. For denervation, a segment of around 5mm of sciatic nerve was surgically removed, as previously described (*78*). Upon denervation, muscle mass variation (%) was calculated as the difference of the mass of the denervated and innervated muscles (from the contralateral leg), normalized to the mass of the innervated muscle. AAV9 carrying shRNA against mouse *Ctbp1*, *Ctbp2* or scramble (Vector Biolabs) were injected in the anterior hind limb compartment (8.10^10^ virus particle). Mice were sacrificed 4 weeks after AAV injection. All animal studies were performed in accordance with the European Union guidelines for animal care and approved by the Veterinary Office of the Canton of Geneva (application number GE220/GE227).

### Cell culture

C2C12 cell line was acquired from ATCC (CRL-1772) and cultured in Dulbecco’s modified Eagle’s medium (DMEM – Sigma D5796) supplemented with 10% fetal bovine serum and 1% Penicillin-Streptomycin (Pen/Strp). Cell differentiation was induced by switching to DMEM with 2% horse serum and 1% Pen/Strep.

### Transcript expression analyses

Total RNA was extracted from TA muscle or cells using RNeasy Fibrous Tissue Mini Kit (Qiagen). DNAse-treated mRNA were reverse transcribed using High-Capacity cDNA Reverse Transcription Kit (ThermoFisher). Quantitative PCR were performed with Power Up Sybr Master Mix (ThermoFisher), analyzed with StepOne software, and normalized on *Tbp* expression. Primers are listed in Supplementary Table 1. For RNAseq, RNA quantification was performed with a Qubit fluorometer (ThermoFisher) and RNA integrity assessed with a Bioanalyzer (Agilent Technologies). RNA-seq libraries were prepared with TruSeq mRNA stranded kit and sequenced on the Illumina HiSeq 4000 sequencer according to the manufacturer’s instruction at the iGE3 platform (Geneva, Switzerland).

Quality control of the dataset was checked using FastQC Version 0.11.9. The FASTQ files will be deposited at Gene Expression Omnibus upon acceptance. Sequence reads were trimmed for adaptor sequence using Trimmomatic, aligned to *M. musculus* genome GRCm39 in R, and transformed to count tables using Rsubread V.2.12.2 package from Bioconductor. Count data were used for differential expression gene (DEG) analysis using Limma V.3.54.1 package and edgR V.3.40.2 from Bioconductor. DEGs between two conditions were filtered based on log2 fold change up or down ≥ 1.5 and statistical p-value < 0.05. PCA plot was generated using R packages (ggfortify, ggplot2, and stats). Venn diagram was created using Venny 2.1.0. Gene ontology analyses were performed using filtered DEG sets against the GeneOntology database using Fgsea V.1.24.0 from Bioconductor packages in R. Dot plots and Volcano plots were generated using the ggplot2 Version 3.4.0 package in R, and heatmaps for individual genes lists were generated using GraphPad Prism Version 9.

### Western blot analysis

Proteins were extracted from nitrogen-powdered TA muscle and cells in RIPA lysis buffer (50mM Tris HCl pH8, 150mM NaCl, 1% NP-40, 0.5% Sodium Deoxycholate, 0.1% SDS, 1% Triton X100, 10% Glycerol) supplemented with proteases (Pierce) and phosphatases inhibitors (Roche). Lysates were sonicated twice 10 seconds, centrifuged at 10’000 RPM for 10min at 4°C, dosed with the BCA Protein Assay (Pierce), separated in polyacrylamide SDS gels, and transferred to nitrocellulose membrane. Membranes were blocked in TBS, 0.1% Tween 20, 3% BSA, and incubated successively with primary antibodies and HRP-coupled secondary antibodies. Proteins were revealed with LumiGLO Peroxidase Chemiluminescent Substrate Kit (Sera Care) and analyzed with the iBright1500 (Invitrogen).

### Inorganic staining and immunostaining

For histology, TA muscles were frozen in nitrogen-cooled isopentane. 8 µm sections were stained with Hematein/Eosin and for NADH dehydrogenase activity (NADH tetrazolium reductase reaction, NADH-TR) as previously described (*79, 80*). Pictures were captured with a widefield Zeiss Axioskop 2 plus microscope. For immunostaining, TA sections and EDL muscles were fixed in 4% paraformaldehyde (PFA) and quenched in PBS, 0.1M Glycine. TA sections were blocked in PBS, 3% IgG free BSA (Jackson Immunoresearch), 0.2% Triton X100. For fiber type staining, TA sections were unfixed and blocked in PBS, 3% IgG free BSA, with AffiniPure Mouse IgG Fab Fragments (Jackson ImmunoResearch). EDL muscles were micro-dissected into bundles or single fibers and blocked in PSB, 3% BSA, 2% Triton X100. Sections, bundles and single fibers were incubated sequentially with primary and fluorescent secondary antibodies, then mounted in Vectashield with or without DAPI (Vector Laboratories). Pictures were acquired using fluorescent widefield Zeiss Axio Imager M2, confocal Axio Imager Z2 Basis LSM 800 or Leica TCS SP8 STED 3X microscope. Fiber type and size distribution was determined based on the laminin staining with Fiji, as previously described (*81*).

### AChRs turnover assay

AChR turnover was assessed by sequential injection of 25 pmoles of α-bungarotoxin-Alexa488 and - Alexa555 (Invitrogen), as previously done (*24*). Pictures of whole-mount EDL muscle fiber bundles were acquired with a confocal Axio Imager Z2 Basis LSM 800. Pixel dominance corresponding to “old” and new AChRs and endplate volume were calculated using Fiji and Matlab software.

### Antibodies

The following antibodies were used: CtBP1 (PA582651; 1/2000 for IF on sections, 1/250 for IF on fibers) from ThermoFisher; Pak1 (2602), CtBP1 (8684; 1/200 for IF) from Cell Signaling; Actinin (A7732) from Sigma; CtBP1 (612042; 1/500 for IF) from BD Biosciences; Tubulin (15246), OXPHOS kit (110413), Laminin (11575; 1/500 for IF) and TOMM20 (186735; 1/250 for IF) from Abcam; Synaptophysin (GT2589; 1/250 for IF) from Genetex Lucerna; Myosin Heavy Chain IIB (BF-F3; 1/25 for IF), IIA (sc-71; 1/50 for IF) from Developmental Studies Hybridoma Bank. All antibodies were diluted at 1/1000 for Western blot.

### Images and Statistical analyses

Images were analyzed with Zen lite, LAS X, Fiji, Imaris and Matlab software. Results are expressed as mean ± SEM of independent samples, with *n* (number of individual experiments) ≥ 3. Statistical comparison was performed using two-tailed Student’s *t*-test, one/two-way ANOVA test, or Kolmogorov-Smirnov test, dependent on the conditions. A 0.05 level of confidence was accepted for statistical significance.

## Acknowledgements

We thank Dr. François Prodon for his help with STED microscope. This work was funded by the Swiss National Science Foundation. OC received funding from the Institute of Genetics and Genomics of Geneva.

## Competing interests

The authors declare no competing interests.

## Notes

### Competing Interest Statement

The authors have declared no competing interest.

## References

1. A. M. Simon, P. Hoppe, S. J. Burden, Spatial restriction of AChR gene expression to subsynaptic nuclei. Development 114, 545–553 (1992).

2. V. Witzemann, H. R. Brenner, B. Sakmann, Neural factors regulate AChR subunit mRNAs at rat neuromuscular synapses. J Cell Biol 114, 125–141 (1991).

3. H. Tang, D. Goldman, Activity-dependent gene regulation in skeletal muscle is mediated by a histone deacetylase (HDAC)-Dach2-myogenin signal transduction cascade. PNAS 103, 16977–16982 (2006).

4. H. Tang et al., CaM kinase II-dependent phosphorylation of myogenin contributes to activity-dependent suppression of nAChR gene expression in developing rat myotubes. Cell Signal 16, 551–563 (2004).

5. A. Mejat et al., Histone deacetylase 9 couples neuronal activity to muscle chromatin acetylation and gene expression. Nat Neurosci 8, 313–321 (2005).

6. L. Berghella et al., A highly conserved molecular switch binds MSY-3 to regulate myogenin repression in postnatal muscle. Genes Dev 22, 2125–2138 (2008).

7. J. L. Thomas et al., PAK1 and CtBP1 Regulate the Coupling of Neuronal Activity to Muscle Chromatin and Gene Expression. Mol Cell Biol 35, 4110–4120 (2015).

8. T. J. Cohen et al., The histone deacetylase HDAC4 connects neural activity to muscle transcriptional reprogramming. J Biol Chem 282, 33752–33759 (2007).

9. T. J. Cohen et al., The deacetylase HDAC4 controls myocyte enhancing factor-2-dependent structural gene expression in response to neural activity. FASEB J. 23, 99–106 (2009).

10. H. Tang et al., A histone deacetylase 4/myogenin positive feedback loop coordinates denervation-dependent gene induction and suppression. Mol Cell Biol 20, 1120–1131 (2009).

11. M. C. Choi et al., A direct HDAC4-MAP kinase crosstalk activates muscle atrophy program. Mol cell 47, 122–132 (2012).

12. V. Moresi et al., Myogenin and class II HDACs control neurogenic muscle atrophy by inducing E3 ubiquitin ligases. Cell 143, 35–45 (2010).

13. M. Kuppuswamy et al., Role of the PLDLS-binding cleft region of CtBP1 in recruitment of core and auxiliary components of the corepressor complex. Mol Cell Biol 28, 269–281 (2008).

14. R. Weigert et al., CtBP/BARS induces fission of Golgi membranes by acylating lysophosphatidic acid. Nature 402, 429–433 (1999).

15. A. Pagliuso et al., Golgi membrane fission requires the CtBP1-S/BARS-induced activation of lysophosphatidic acid acyltransferase delta. Nat Commun 7, 12148 (2016).

16. M. Bonazzi et al., CtBP3/BARS drives membrane fission in dynamin-independent transport pathways. Nat Cell Biol 7, 570–580 (2005).

17. C. J. Barnes et al., Functional inactivation of a transcriptional corepressor by a signaling kinase. Nat Struct Biol 10, 622–628 (2003).

18. D. Ivanova et al., CtBP1-Mediated Membrane Fission Contributes to Effective Recycling of Synaptic Vesicles. Cell Rep 30, 2444–2459 e2447 (2020).

19. H. Jafari Khamirani et al., Exome sequencing identified a de novo frameshift pathogenic variant of CTBP1 in an extremely rare case of HADDTS. J Genet 100, (2021).

20. E. W. Sommerville et al., De novo CTBP1 variant is associated with decreased mitochondrial respiratory chain activities. Neurol Genet 3, e187 (2017).

21. W. K. Wong et al., Mitochondrial respiratory chain dysfunction in a patient with a heterozygous de novo CTBP1 variant. JIMD Rep 63, 546–554 (2022).

22. M. A. Blevins, M. Huang, R. Zhao, The Role of CtBP1 in Oncogenic Processes and Its Potential as a Therapeutic Target. Mol Cancer Ther 16, 981–990 (2017).

23. D. Ivanova et al., Synaptic activity controls localization and function of CtBP1 via binding to Bassoon and Piccolo. EMBO J 34, 1056–1077 (2015).

24. P. Castets et al., mTORC1 and PKB/Akt control the muscle response to denervation by regulating autophagy and HDAC4. Nat Commun 10, 3187 (2019).

25. I. Martinez-Penay Valenzuela, C. Mouslim, M. Akaaboune, Calcium/calmodulin kinase II-dependent acetylcholine receptor cycling at the mammalian neuromuscular junction in vivo. J Neurosci 30, 12455–12465 (2010).

26. S. Strack et al., A Novel Labeling Approach Identifies Three Stability Levels of Acetylcholine Receptors in the Mouse Neuromuscular Junction In Vivo. PLoS One 6, e20524 (2011).

27. J. H. Kim, H. D. Youn, C-terminal binding protein maintains mitochondrial activities. Cell Death Differ 16, 584–592 (2009).

28. M. Grooteclaes et al., C-terminal-binding protein corepresses epithelial and proapoptotic gene expression programs. Proc Natl Acad Sci U S A 100, 4568–4573 (2003).

29. O. Liska et al., TFLink: an integrated gateway to access transcription factor-target gene interactions for multiple species. Database (Oxford) 2022, (2022).

30. L. Q. Lou, W. Q. Zhou, X. Song, Z. Chen, Elevation of hsa-miR-7-5p level mediated by CtBP1-p300-AP1 complex targets ATXN1 to trigger NF-kappaB-dependent inflammation response. J Mol Med (Berl) 101, 223–235 (2023).

31. L. Bai et al., CtBP proteins transactivate matrix metalloproteinases and proinflammatory cytokines to mediate the pathogenesis of abdominal aortic aneurysm. Exp Cell Res 421, 113386 (2022).

32. D. Deutsch et al., Tuftelin: enamel mineralization and amelogenesis imperfecta. Ciba Found Symp 205, 135–147; discussion 147-155 (1997).

33. B. Zhu, M. Zhang, S. D. Byrum, A. J. Tackett, J. K. Davie, TBX2 blocks myogenesis and promotes proliferation in rhabdomyosarcoma cells. Int J Cancer 135, 785–797 (2014).

34. C. Li, X. Q. Xiao, Y. H. Qian, Z. Y. Zhou, The CtBP1-p300-FOXO3a transcriptional complex represses the expression of the apoptotic regulators Bax and Bim in human osteosarcoma cells. J Cell Physiol 234, 22365–22377 (2019).

35. H. Tang et al., mTORC1 promotes denervation-induced muscle atrophy through a mechanism involving the activation of FoxO and E3 ubiquitin ligases. Sci. Signal. 7, ra18 (2014).

36. C. Ninfali, L. Siles, D. S. Darling, A. Postigo, Regulation of muscle atrophy-related genes by the opposing transcriptional activities of ZEB1/CtBP and FOXO3. Nucleic Acids Res 46, 10697–10708 (2018).

37. P. Nagpal et al., The ubiquitin ligase Nedd4-1 participates in denervation-induced skeletal muscle atrophy in mice. PLoS One 7, e46427 (2012).

38. J. Shi, L. Luo, J. Eash, C. Ibebunjo, D. J. Glass, The SCF-Fbxo40 complex induces IRS1 ubiquitination in skeletal muscle, limiting IGF1 signaling. Dev Cell 21, 835–847 (2011).

39. X. Lin et al., Opposed regulation of corepressor CtBP by SUMOylation and PDZ binding. Mol Cell 11, 1389–1396 (2003).

40. M. H. Kagey, T. A. Melhuish, D. Wotton, The polycomb protein Pc2 is a SUMO E3. Cell 113, 127–137 (2003).

41. Y. Izumiya et al., Fast/Glycolytic muscle fiber growth reduces fat mass and improves metabolic parameters in obese mice. Cell Metab 7, 159–172 (2008).

42. O. Kirsh et al., The SUMO E3 ligase RanBP2 promotes modification of the HDAC4 deacetylase. EMBO J 21, 2682–2691 (2002).

43. S. Gregoire, X. J. Yang, Association with class IIa histone deacetylases upregulates the sumoylation of MEF2 transcription factors. Mol Cell Biol 25, 2273–2287 (2005).

44. C. Riquelme, K. K. Barthel, X. Liu, SUMO-1 modification of MEF2A regulates its transcriptional activity. J Cell Mol Med 10, 132–144 (2006).

45. C. L. Zhang, T. A. McKinsey, J. R. Lu, E. N. Olson, Association of COOH-terminal-binding protein (CtBP) and MEF2-interacting transcription repressor (MITR) contributes to transcriptional repression of the MEF2 transcription factor. J Biol Chem 276, 35–39 (2001).

46. L. Siles et al., ZEB1 imposes a temporary stage-dependent inhibition of muscle gene expression and differentiation via CtBP-mediated transcriptional repression. Mol Cell Biol 33, 1368–1382 (2013).

47. U. Dressel et al., A dynamic role for HDAC7 in MEF2-mediated muscle differentiation. J Biol Chem 276, 17007–17013 (2001).

48. A. Ravel-Chapuis, M. Vandromme, J. L. Thomas, L. Schaeffer, Postsynaptic chromatin is under neural control at the neuromuscular junction. The EMBO journal 26, 1117–1128 (2007).

49. I. Durr, M. Numberger, C. Berberich, V. Witzemann, Characterization of the functional role of E-box elements for the transcriptional activity of rat acetylcholine receptor epsilon-subunit and gamma-subunit gene promoters in primary muscle cell cultures. Eur J Biochem 224, 353–364 (1994).

50. X. Chen et al., The CtIP-CtBP1/2-HDAC1-AP1 transcriptional complex is required for the transrepression of DNA damage modulators in the pathogenesis of osteosarcoma. Transl Oncol 21, 101429 (2022).

51. J. H. Kim, E. J. Cho, S. T. Kim, H. D. Youn, CtBP represses p300-mediated transcriptional activation by direct association with its bromodomain. Nat Struct Mol Biol 12, 423–428 (2005).

52. C. C. Fjeld, W. T. Birdsong, R. H. Goodman, Differential binding of NAD+ and NADH allows the transcriptional corepressor carboxyl-terminal binding protein to serve as a metabolic sensor. Proc Natl Acad Sci U S A 100, 9202–9207 (2003).

53. V. Kumar et al., Transcription corepressor CtBP is an NAD(+)-regulated dehydrogenase. Mol Cell 10, 857–869 (2002).

54. Q. Zhang, D. W. Piston, R. H. Goodman, Regulation of corepressor function by nuclear NADH. Science 295, 1895–1897 (2002).

55. D. L. Madison, J. A. Wirz, D. Siess, J. R. Lundblad, Nicotinamide adenine dinucleotide-induced multimerization of the co-repressor CtBP1 relies on a switching tryptophan. J Biol Chem 288, 27836–27848 (2013).

56. A. G. Bellesis, A. M. Jecrois, J. A. Hayes, C. A. Schiffer, W. E. Royer, Jr., Assembly of human C-terminal binding protein (CtBP) into tetramers. J Biol Chem 293, 9101–9112 (2018).

57. A. Verger et al., Mechanisms directing the nuclear localization of the CtBP family proteins. Mol Cell Biol 26, 4882–4894 (2006).

58. J. C. Nichols, C. A. Schiffer, W. E. Royer, Jr., NAD(H) phosphates mediate tetramer assembly of human C-terminal binding protein (CtBP). J Biol Chem 296, 100351 (2021).

59. M. Garriga-Canut et al., 2-Deoxy-D-glucose reduces epilepsy progression by NRSF-CtBP-dependent metabolic regulation of chromatin structure. Nat Neurosci 9, 1382–1387 (2006).

60. Y. Shen et al., Bioenergetic state regulates innate inflammatory responses through the transcriptional co-repressor CtBP. Nat Commun 8, 624 (2017).

61. M. Kreuzer et al., Glycolysis, via NADH-dependent dimerisation of CtBPs, regulates hypoxia-induced expression of CAIX and stem-like breast cancer cell survival. FEBS Lett 594, 2988–3001 (2020).

62. J. S. Elmendorf, A. Damrau-Abney, T. R. Smith, T. S. David, J. Turinsky, Phosphatidylinositol 3-kinase and dynamics of insulin resistance in denervated slow and fast muscles in vivo. Am J Physiol 272, E661–670 (1997).

63. J. Turinsky, Dynamics of insulin resistance in denervated slow and fast muscles in vivo. The Am J Physiol 252, R531–537 (1987).

64. J. P. Jones, E. B. Tapscott, A. L. Olson, J. E. Pessin, G. L. Dohm, Regulation of glucose transporters GLUT-4 and GLUT-1 gene transcription in denervated skeletal muscle. J Appl Physiol (1985) 84, 1661–1666 (1998).

65. A. Handberg et al., Reciprocal GLUT-1 and GLUT-4 expression and glucose transport in denervated muscles. Am J Physiol 271, E50–57 (1996).

66. L. Coderre et al., Alteration in the expression of GLUT-1 and GLUT-4 protein and messenger RNA levels in denervated rat muscles. Endocrinology 131, 1821–1825 (1992).

67. A. Vainshtein, E. M. Desjardins, A. Armani, M. Sandri, D. A. Hood, PGC-1alpha modulates denervation-induced mitophagy in skeletal muscle. Skeletal muscle 5, 9 (2015).

68. S. M. Hughes, M. M. Chi, O. H. Lowry, K. Gundersen, Myogenin induces a shift of enzyme activity from glycolytic to oxidative metabolism in muscles of transgenic mice. J Cell Biol 145, 633–642 (1999).

69. M. Triolo, M. Slavin, N. Moradi, D. A. Hood, Time-dependent changes in autophagy, mitophagy and lysosomes in skeletal muscle during denervation-induced disuse. J Physiol 600, 1683–1701 (2022).

70. Y. T. Kuo, P. H. Shih, S. H. Kao, G. C. Yeh, H. M. Lee, Pyrroloquinoline Quinone Resists Denervation-Induced Skeletal Muscle Atrophy by Activating PGC-1alpha and Integrating Mitochondrial Electron Transport Chain Complexes. PLoS One 10, e0143600 (2015).

71. P. J. Adhihetty, M. F. O’Leary, B. Chabi, K. L. Wicks, D. A. Hood, Effect of denervation on mitochondrially mediated apoptosis in skeletal muscle. J Appl Physiol (1985) 102, 1143–1151 (2007).

72. K. Singh, D. A. Hood, Effect of denervation-induced muscle disuse on mitochondrial protein import. Am J Physiol Cell Physiol 300, C138–145 (2011).

73. A. Wagatsuma, N. Kotake, K. Mabuchi, S. Yamada, Expression of nuclear-encoded genes involved in mitochondrial biogenesis and dynamics in experimentally denervated muscle. J Physiol Biochem 67, 359–370 (2011).

74. J. M. Memme, M. Slavin, N. Moradi, D. A. Hood, Mitochondrial Bioenergetics and Turnover during Chronic Muscle Disuse. Int J Mol Sci 22, (2021).

75. T. Yokokawa et al., Muscle denervation reduces mitochondrial biogenesis and mitochondrial translation factor expression in mice. Biochem Biophys Res Commun 527, 146–152 (2020).

76. M. Joffe, N. Savage, H. Isaacs, Biochemical functioning of mitochondria in normal and denervated mammalian skeletal muscle. Muscle Nerve 4, 514–519 (1981).

77. Y. Kim et al., Reorganization of mitochondria-organelle interactions during postnatal development in skeletal muscle. J Physiol 602, 891–912 (2024).

78. T. Shavlakadze, J. White, J. F. Hoh, N. Rosenthal, M. D. Grounds, Targeted expression of insulin-like growth factor-I reduces early myofiber necrosis in dystrophic mdx mice. Mol Ther 10, 829–843 (2004).

79. V. Dubowitz, C. Sewry, Muscle Biopsy: A Practical Approach. (Elsevier Health Sciences, London, ed. 4th edition, 2013).

80. A. Prola et al., Cardiolipin content controls mitochondrial coupling and energetic efficiency in muscle. Sci Adv 7, (2021).

81. D. J. Ham et al., The neuromuscular junction is a focal point of mTORC1 signaling in sarcopenia. Nat Commun 11, 4510 (2020).

