## Supplemental Data for "CtBP1 coordinates synaptic, metabolic and contractile changes induced by denervation in skeletal muscle"

Supplementary data includes: supplementary references, 9 supplementary figures, 1 supplementary table.

### Suppementary references

1. D. Li, S. Hsu, D. Purushotham, R. L. Sears, T. Wang, WashU Epigenome Browser update 2019. *Nucleic Acids Res.* **47**, W158-W165 (2019).
2. D. Li *et al.*, WashU Epigenome Browser update 2022. *Nucleic Acids Res.* **50**, W774-781 (2022).

### Supplementary Figures

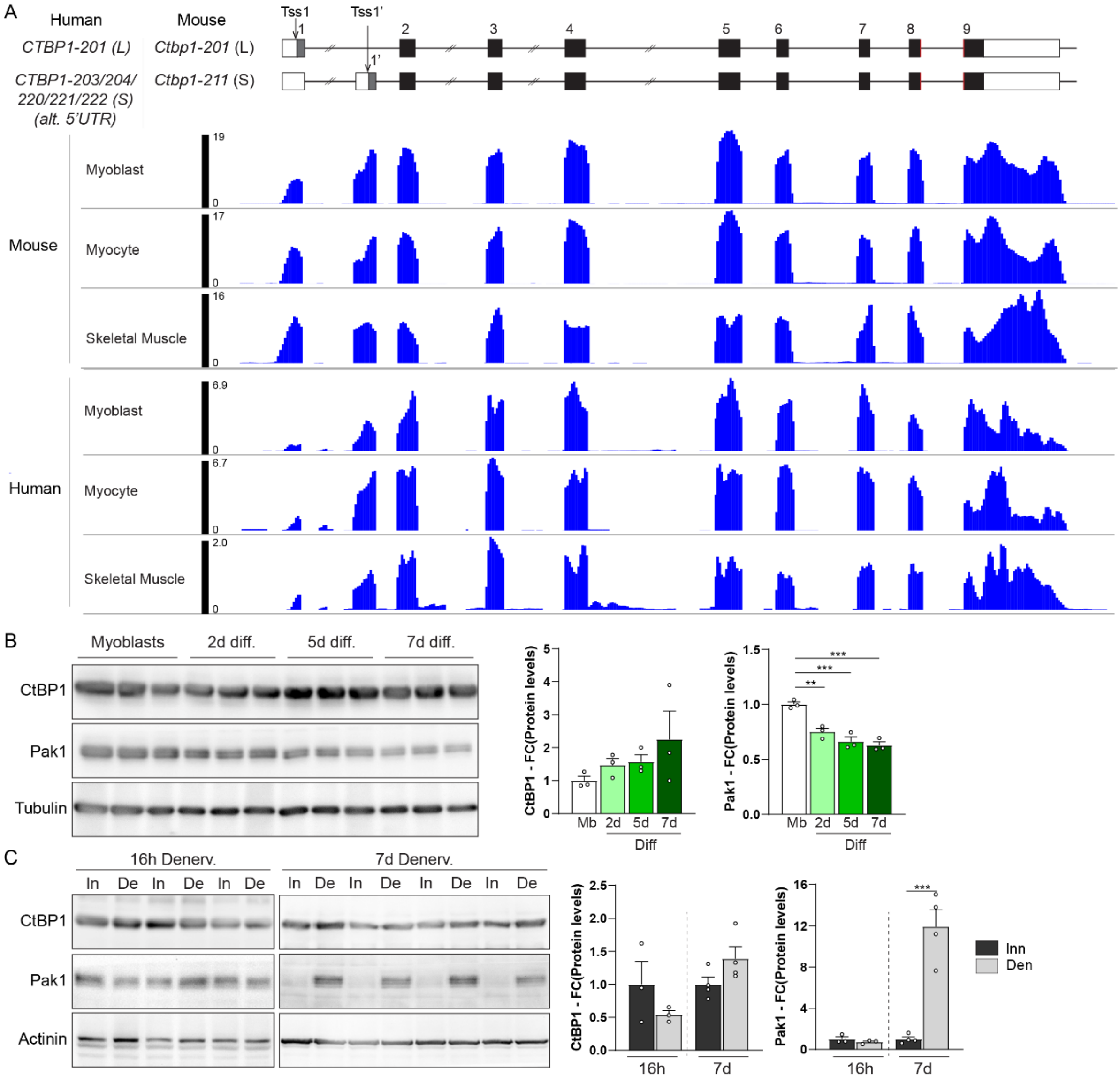


**fig. S1: Expression of CtBP1 isoforms is unchanged upon C2C12 differentiation and muscle denervation.** (**A**) Organization of *Ctbp1/CTBP1* genes in human and mouse. Exons, introns and untranslated (UTR) sequences are represented by grey boxes, black lines and white boxes, respectively. Alternative ATG codons are indicated with arrows. RNAseq data from *WashU Epigenome Browser* (*1*, *2*) confirm the expression of *Ctbp1-l* and *Ctbp1-s* in muscle cells from mouse, while human muscle cells express mainly *CTBP1-S*. (**B**) Western Blot analysis of CtBP1 and Pak1 in C2C12 myoblasts and after 2 to 7 days (d) of differentiation (diff). Protein levels are normalized to tubulin and to myoblasts. (**C**) Western Blot analysis of CtBP1 and Pak1 in innervated muscle (In) and 16h or 7 days (De) after denervation (De). Protein levels are normalized to actinin and to innervated muscle. All values are mean ± s.e.m.; n=3 (B, C for 16h) and 4 (C for 7 days); **$p<0.01$ ***$p<0.001$; one-way ANOVA with Tukey's post-hoc (B) and Student's t-test (C).

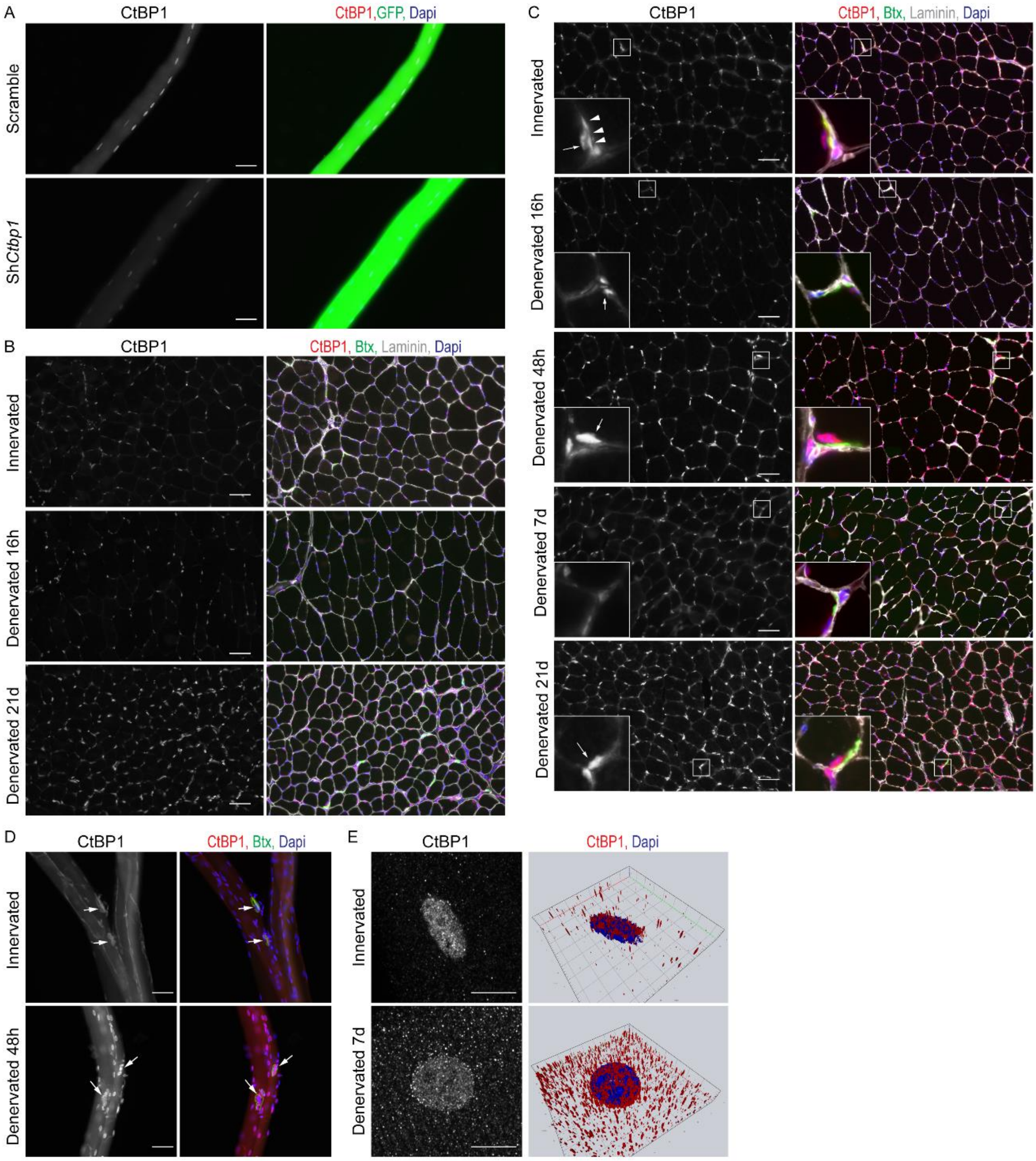


**fig. S2. CtBP1 accumulates in myonuclei 48h after denervation.** (**A**) Staining of single EDL fibers with CtBP1 antibody and GFP reveals after AAV-shCtbp1 infection confirms the specificity of CtBP1 antibody. Scale bar, 50µm. (**B** and **C**) Immunostaining of innervated and denervated (16h to 21 days) TA sections with rabbit (B) and mouse (C) CtBP1 antibody shows transient accumulation of CtBP1 in non- and sub- synaptic myonuclei 48h after nerve injury. Scale bar, 50µm. (**D**) Immunostaining of innervated and denervated (48h) single EDL fibers with mouse CtBP1 antibody shows transient accumulation of CtBP1 in non- and sub- (arrows) synaptic myonuclei 48h after nerve injury. Scale bar, 50µm. (**E**) Enlarged view and 3D reconstruction of nuclear CtBP1 staining in innervated and denervated (7 days) TA muscles. Scale bar, 10µm.

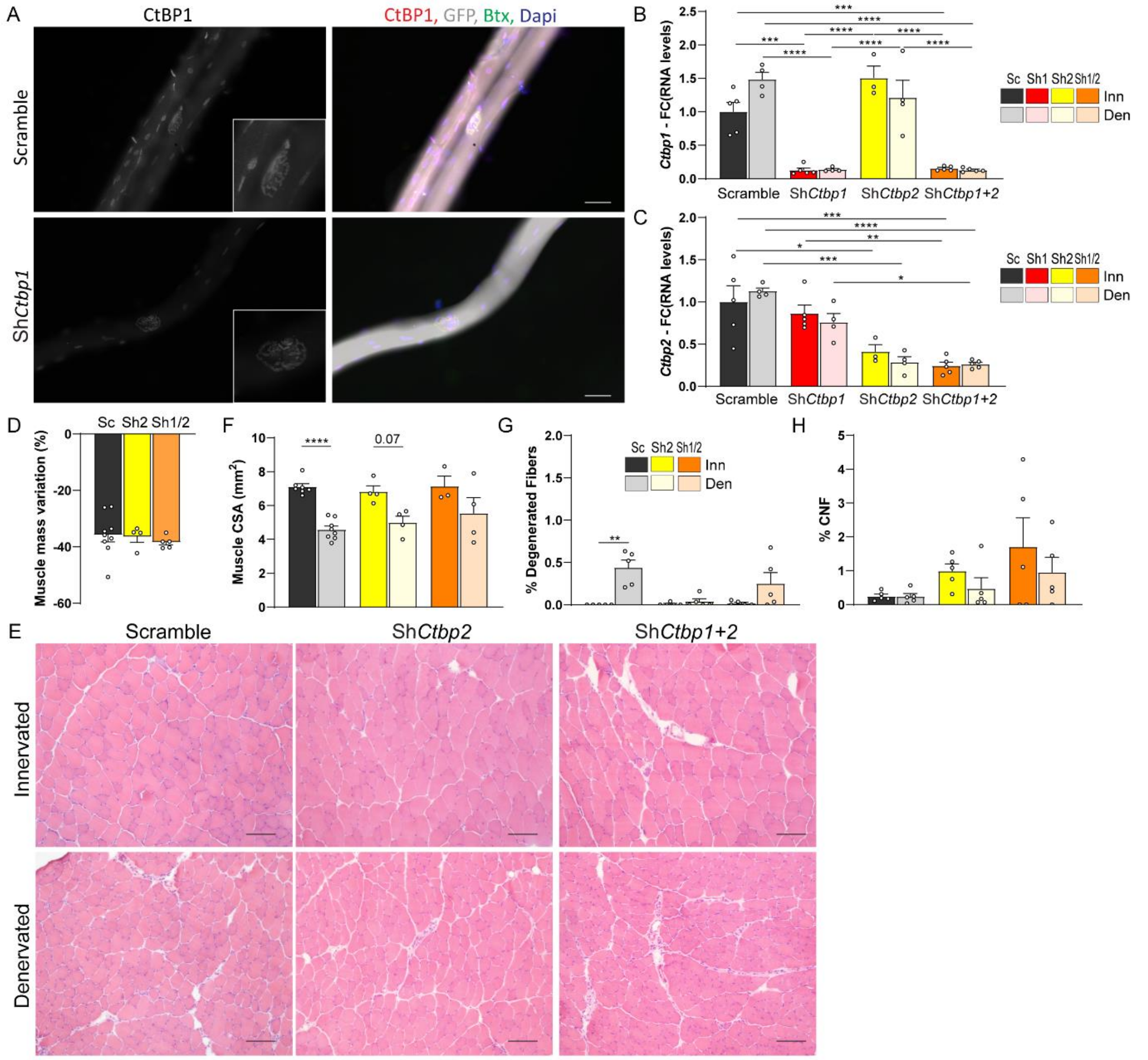

**fig. S3. *Ctbp2* knockdown does not affect Ctbp1 expression or muscle histology.** (**A**) Immunostaining with CtBP1 antibody of innervated single EDL fibers shows loss of CtBP1 staining in the muscle fiber, but not in the pre-synaptic compartment, after infection with AAV-sh*Ctbp1*. Scale bar, 50µm. (**B** and **C**) mRNA levels of *Ctbp1* (B) and *Ctbp*2 (C) in innervated (Inn) and denervated (Den) TA muscles infected with AAV-*shCtbp1* (Sh1), -sh*Ctbp2* (Sh2), -sh*Ctbp1* and –sh*Ctbp2* (Sh1/2), or -shScramble (Sc). Levels are relative to *Tbp* mRNA and to Scramble innervated. (**D**) Mass variation after 2 weeks of denervation for TA muscles injected with AAV-*shCtbp2* (Sh2), AAV-*shCtbp1* and -*shCtbp2* (Sh1/2) or -shScramble (Sc). (**E**) H&E staining of innervated and 2-week-denervated TA muscles injected with AAV-*shCtbp2*, or AAV-*shCtbp1* and -*shCtbp2*, or -shScramble. Scale bar, 100µm. (**F**) Cross sectional area (CSA) of innervated and denervated (2 weeks) TA muscles injected with AAV-*shCtbp2* (Sh2), AAV-*shCtbp1* and -*shCtbp2* (Sh1/2) or -shScramble (Sc). (**G** and **H**) The proportion of degenerated fibers (G) and of centronucleated fibers (CNF, H) is unchanged in innervated and denervated TA muscles injected with AAV-*shCtbp2* (Sh2), AAV-*shCtbp1* and -*shCtbp2* (Sh1/2), compared to Scramble (Scr). All values are mean ± s.e.m.; n=5Sc/Sh1/Sh1+2 (Inn); 4Sc/Sh1/Sh2 (Den); 3InSh2; 5Sh1+2Den (B, C), 4Sh2/5Sh1+2 (D), 4 (F, except Sh1+2Inn, n=3), 5 (G, H); *p<0.05, **p<0.01, ***p<0.001, ****p<0.0001; two-way ANOVA with Tukey's post-hoc.

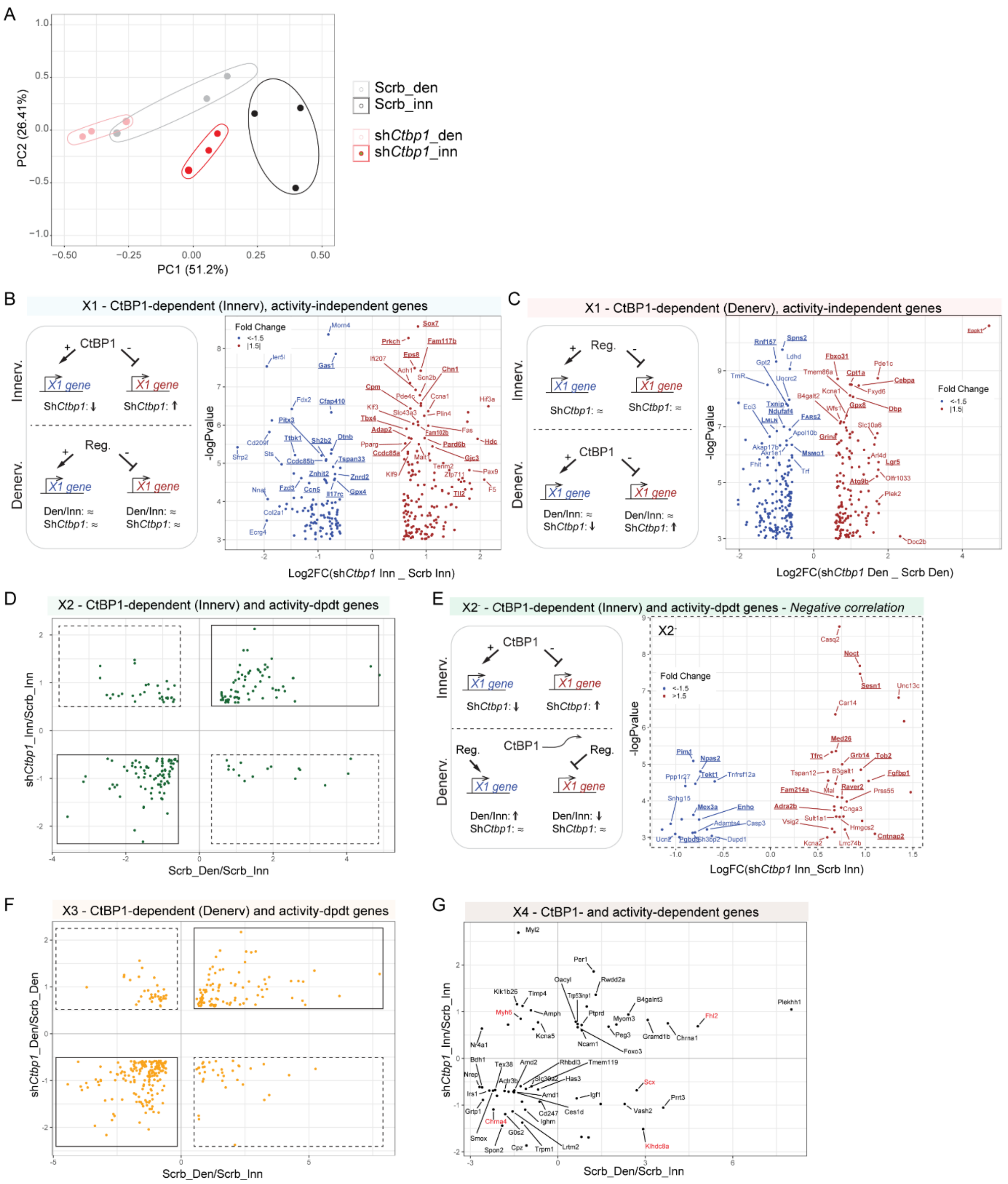


**fig. S4. Transcriptomic analyses of CtBP1-deficient TA muscle 48h after denervation.** (**A**) Principal component analysis (PCA) showing separation of innervated (Inn) vs. denervated (Den), and AAV-sh*Ctbp1* vs. –shScramble groups. (**B** and **C**) *Ctbp1* knockdown alters the expression of activity-independent X1 genes in innervated (B) or denervated (C) muscles. Volcano plots show genes DE in AAV-sh*Ctbp1* innervated (B) or denervated (C) muscles (*vs*. AAV-Scrb). (**D** and **E**) *Ctbp1* knockdown affects the expression of activity-dependent X2 genes only in innervated muscle, with positive (full line) or negative (dotted line) correlation with the denervation effect. The volcano plot in E shows genes DE in AAV-sh*Ctbp1* innervated muscles (*vs*. AAV-Scrb innervated), with negative correlation with denervation-induced changes. (**F**) *Ctbp1* knockdown affects the expression of activity-dependent X3 genes only in denervated muscle, with positive (full line) or negative (dotted line) correlation with the denervation effect. (**G**) Scatter plot showing activity-dependent genes DE in both AAV-sh*Ctbp1* innervated muscles (*vs*. AAV-Scrb innervated) and AAV-sh*Ctbp1* denervated muscles (*vs*. AAV-Scrb denervated). The plot shows the correlation between the effect of denervation and of AAV-sh*Ctbp1* in innervated muscle. Only some X4 genes (red) show an inverse correlation in denervated muscle (see Fig. S5G).

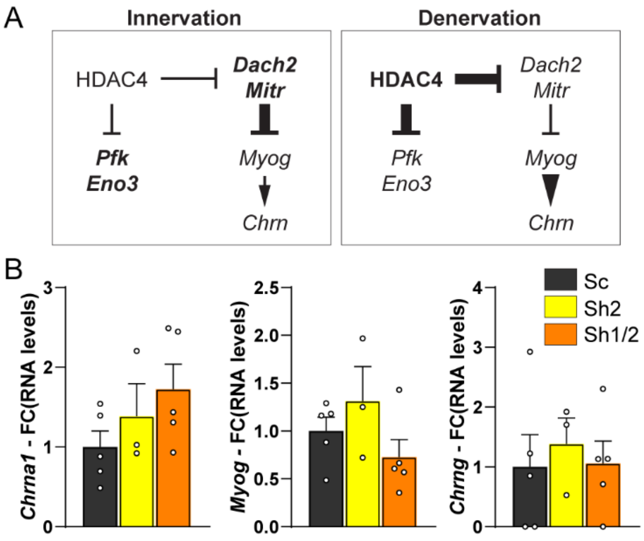


**fig. S5. *Ctbp2* knockdown does not alter synaptic gene expression.** (**A**) HDAC4-dependent regulation of metabolic and synaptic genes in innervated and denervated conditions. (**B**) mRNA levels of *Chrna1*, *Myog* and *Chrng* after AAV-shScramble (Sc), -sh*CtBP2* (Sh2), or -sh*Ctbp1* and -sh*Ctbp2* (Sh1/2) injection in innervated or denervated TA muscles. Levels are relative to *Tbp* mRNA and normalized to AAV-shScramble innervated muscle. All values are mean ± s.e.m.; n=5Sc/Sh1/2, 3Sh2 (B).

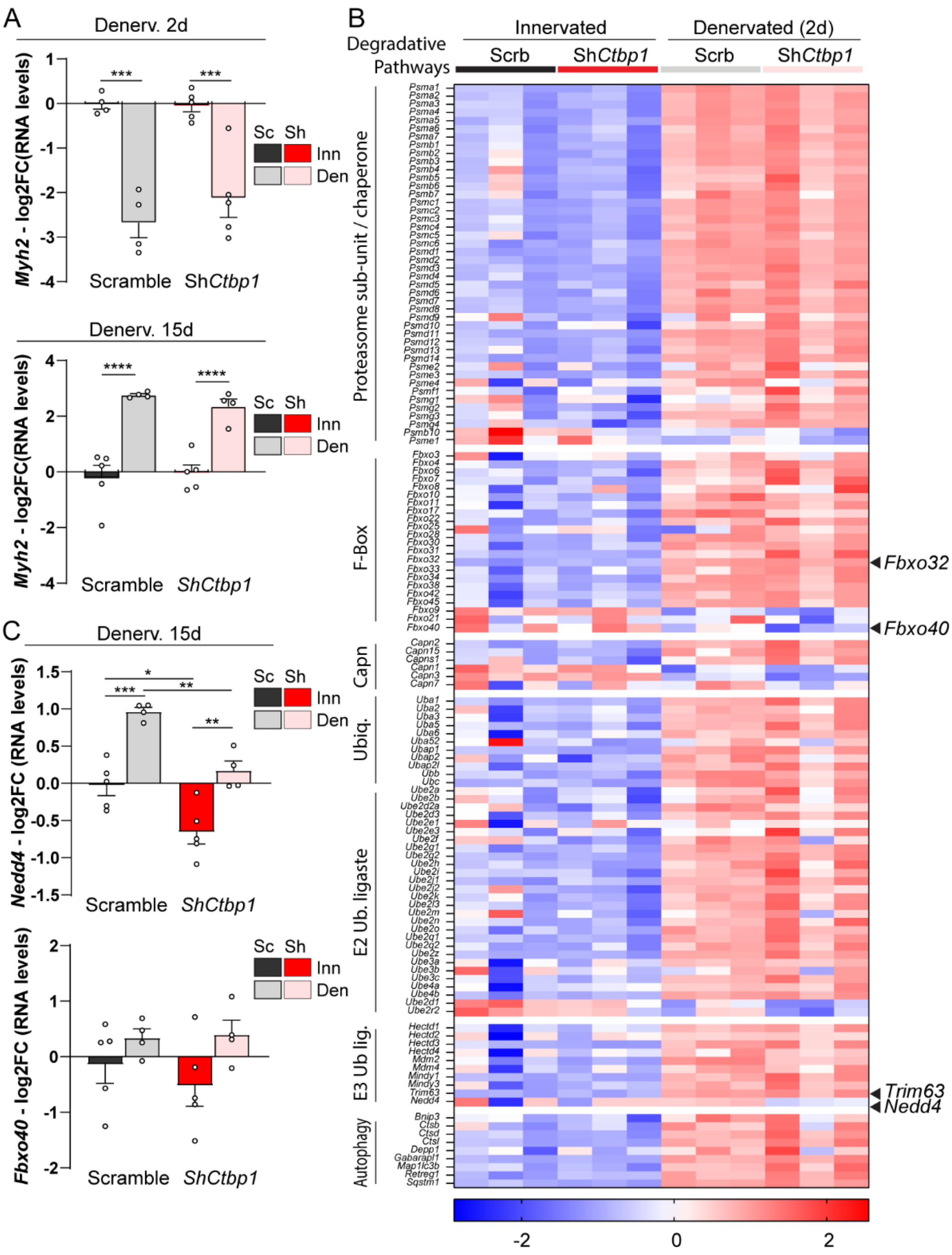


**fig. S6. *Ctbp1* knockdown does not affect genes linked to muscle atrophy.** (**A**) mRNA levels of *Myh2,* encoding myosin heavy chains IIA after AAV-sh*Ctbp1* (Sh) or -shScramble (Sc) injection in innervated (Inn) and 2-day- and 2-week-denervated (Den) TA muscles. Levels are relative to *Tbp* mRNA and normalized to AAV-shScramble innervated muscle. (**B**) Heatmap of z-scores computed based on log2FC of RNAseq counts for genes encoding proteins involved in proteolysis, in innervated and denervated, AAV-sh*Ctbp1* and -shScramble muscles. (**C**) mRNA levels of *Nedd4* and *Fbxo40* after AAV-sh*Ctbp1* (Sh) or -shScramble (Sc) injection in innervated (Inn) and 2-week-denervated (Den) TA muscles. Levels are relative to *Tbp* mRNA and normalized to AAV-shScramble innervated muscle. All values are mean ± s.e.m.; n=4Sc/5Sh (A, 2d); 5Inn/4Den (A, 15d and C); *p<0.05, **p<0.01, ***p<0.001, ****p<0.0001; two- way ANOVA with Tukey’s post-hoc.

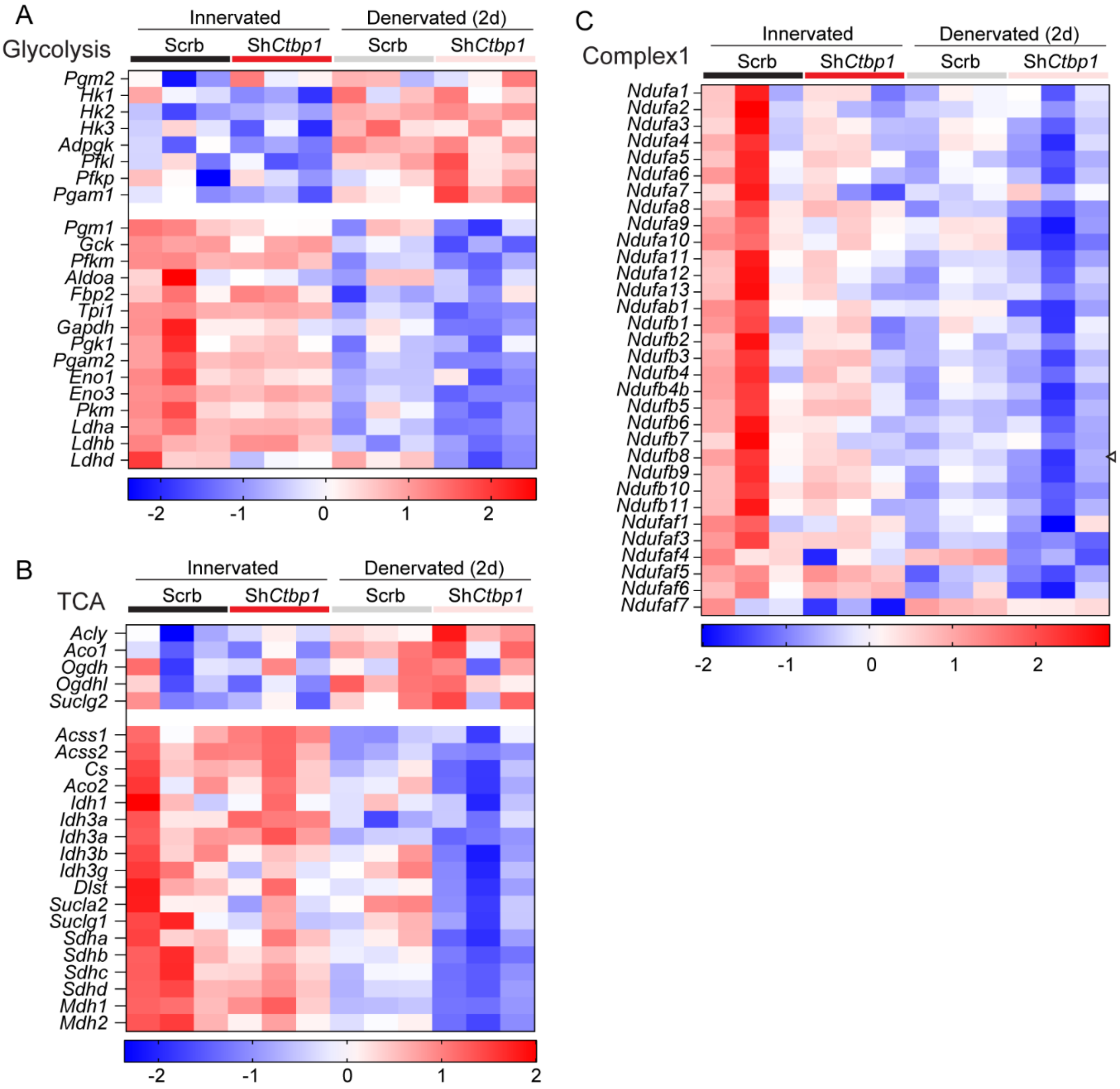


**fig. S7. *Ctbp1* knockdown exacerbates the effect of denervation on metabolic genes.** (**A-C**) Heatmap of z-scores computed based on log2FC of RNAseq counts for nuclear genes encoding proteins involved in glycolysis (A), in tricarboxylic acid (TCA) cycle (B), and in respiratory chain complexes I (C), in innervated and denervated, AAV-sh*Ctbp1* and -shScramble muscles.

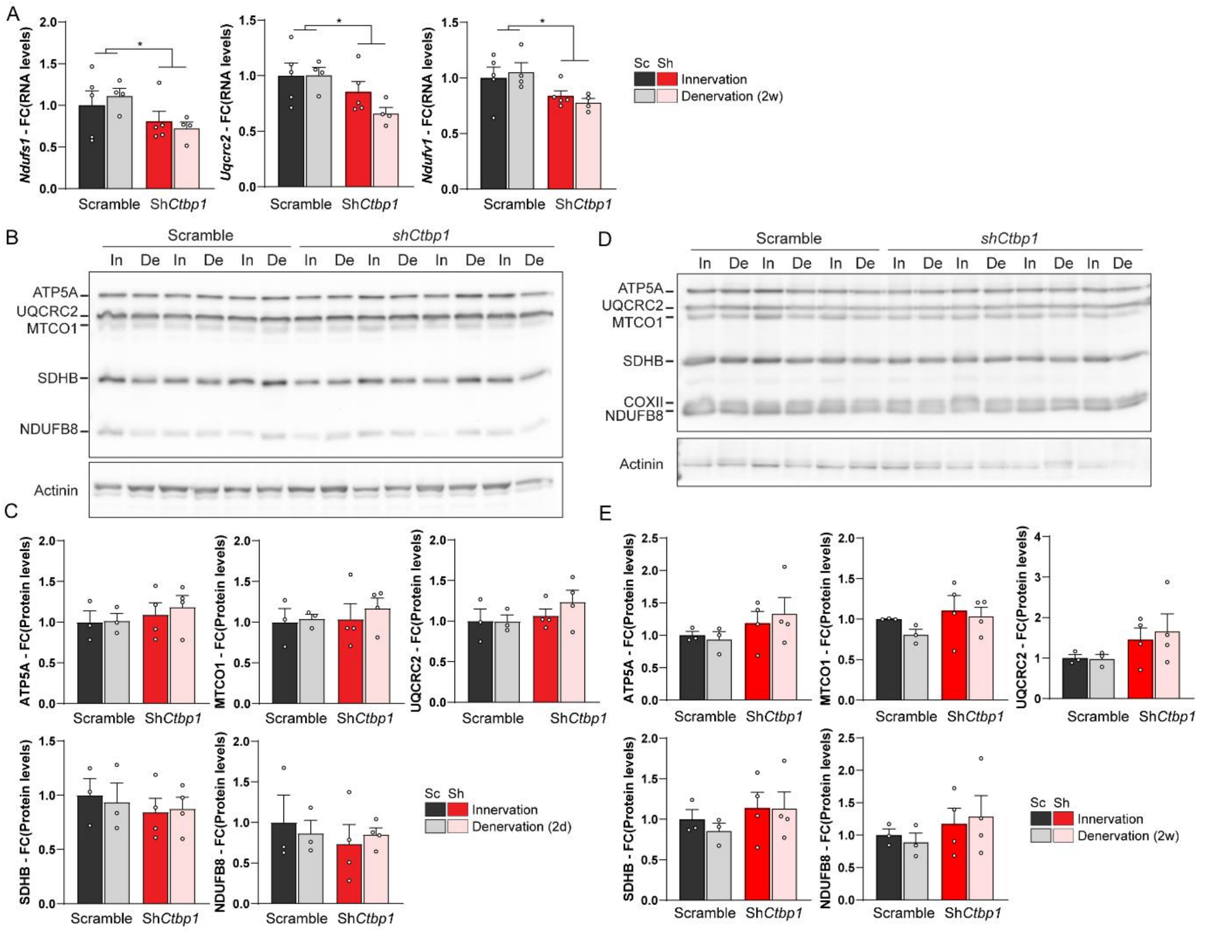


**fig. 8. *Ctbp1* knockdown does not change protein levels of respiratory chain complexes.** (**A**) mRNA levels of genes encoding components of the respiratory complex I (*Ndufs1*, *Ndufv1*) and III (*Uqcrc2*), after AAV-sh*Ctbp1* (Sh) or -shScramble (Sc) injection in innervated (Inn) and 2-week-denervated (Den) TA muscles. Levels are relative to *Tbp* mRNA and normalized to AAV-shScramble innervated muscle. (**B-E**) Protein levels of OXPHOS proteins from innervated (In) and 2-day- (B, C) or 2-week- (D, E) denervated (De) TA muscles infected with AAV-sh*Ctbp1* (Sh) or –shScramble (Sc). Protein levels are normalized to actinin and relative to Scramble innervated. All values are mean ± s.e.m.; n=5Inn/4Den (A), 3Sc/4Sh (B-E); *p<0.05; two-way ANOVA with Tukey's post-hoc.

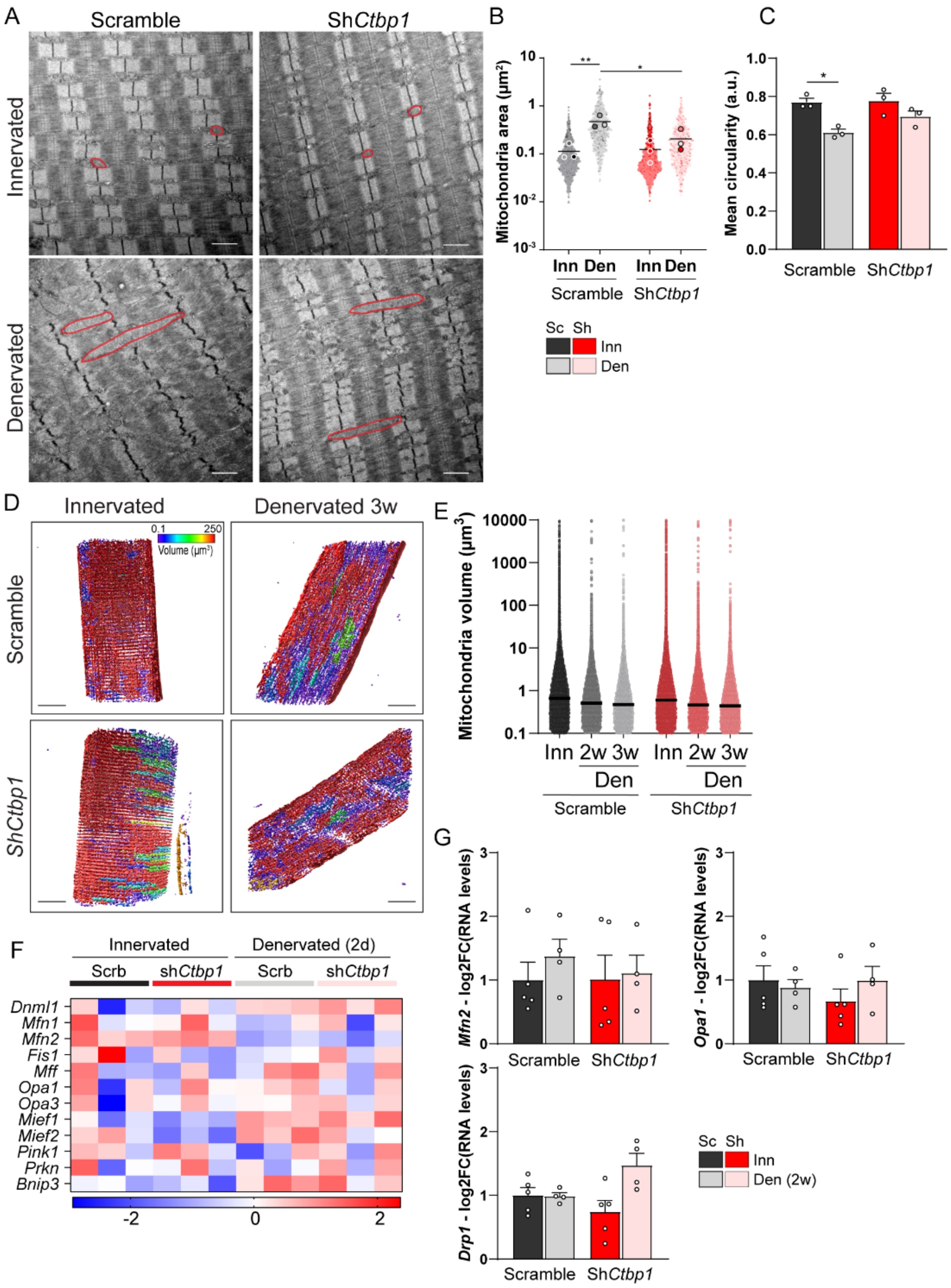


**fig. S9. *Ctbp1* knockdown affects mitochondria network.** (**A - C**) Electron microcopy of innervated and 2-week-denervated TA muscles injected with AAV-sh*Ctbp1* or -shScramble shows elongated mitochondria after denervation. Mitochondria area is given in B; mean circularity of mitochondria is given in C. (**D** and **E**) Imaris 3D reconstitution of mitochondria network in innervated and 3-week-denervated TA muscles injected with AAV-sh*Ctbp1* or -shScramble, with color code corresponding to mitochondria volume. Original pictures are given in Fig. 8E. The distribution of mitochondria volume is given in E. (**F**) Heatmap of z-scores computed based on log2FC of RNAseq counts for genes encoding proteins involved in mitochondria dynamics, in innervated and 2-day-denervated, AAV-sh*Ctbp1* and -shScramble muscles. (**G**) mRNA levels of genes encoding *Mfn2*, *Opa1* and *Drp1* after AAV-sh*Ctbp1* (Sh) or -shScramble (Sc) injection in innervated (Inn) and 2-week-denervated (Den) TA muscles. Levels are relative to *Tbp* mRNA and normalized to AAV-shScramble innervated muscle. All values are mean ± s.e.m.; n=3 (B, C); 5 (E); 5Inn/4Den (G); *p<0.05; **p<0.01; two- way ANOVA with Tukey's post-hoc.

**Supplementary Table1**

| Gene name | Forward primer | Reverse primer |
|---|---|---|
| *Ctbp1-L* | CTGGGCGTCCGACCTCCCATC | TCAGGATAGGCATCTCCACTG |
| *Ctbp1-S* | AATTCATGGTCGTGGAAACC | TCAGGATAGGCATCTCCACTG |
| *Ctbp1-Pan* | AGAGACCTTGGGCATCATTG | CCGCTCGATTCCATCAGATA |
| *Pak1* | ACACGGTTCGAGAAGATTGG | TCCCTCATGACCAGGATCTC |
| *Myog* | CACTCCCTTACGTCCATCGT | CAGGGCTGTTTTCTGGACAT |
| *Chrna1* | TCCCTTCGATGAGCAGAACT | GGGCAGCAGGAGTAGAACAC |
| *Chrng* | GTGTCTTCGAGGTGGCTCTC | ACAGAGATGGAGCAGGAGGA |
| *Chrne* | TTCCCCTTTGACTGGCAGAA | AAAAGCTGCCGTGTCAATGT |
| *Hdac4* | CAGACAGCAAGCCCTCCTAC | AGACCTGTGGTGAACCTTGG |
| *Dach2* | CCAGCTCAAATCCCAGTCAT | CGCAGTTCCTTCTTTTCCTG |
| *Mitr* | CCACCTTGAAGAAGCAGAGG | TGGTGTCCTTAGAGGCTGCT |
| *Pfk* | GATGCAAGGACTTCCGAGAG | GCTCCACTCTGAACGGAAAG |
| *Mse* | GGGAGATGACCTCACGGTAA | TTACAGGCCTGGATGGACTC |
| *Myh2* | ACAAATCTATCCAAGTTCCG | TTCGGTCATTCCACAGCATC |
| *Myl2* | CGTGTTCCTCACGATGTTTG | CCTCTCTGCTTGTGTGGTCA |
| *Myh4* | CAGATGAAAAGGTGGCCATT | CTTCCCTTTGCTTTTGCTTG |
| *Mfn2* | GCACTTTGTCACTGCCAAGA | TGTGTTCCTGTGGGTGTCTT |
| *Opa1* | GAAGGACGACAAAGGCATCC | CCCGTGGTAGGTGATCTTGT |
| *Drp1* | TGCCTCAGATCGTCGTAGTG | GAAACGTGGACTAGCTGCAG |
